# Insect cell plasma membranes do, while soluble enzymes do not, need stabilization by accumulated cryoprotectant molecules during freezing stress

**DOI:** 10.1101/2022.06.23.497306

**Authors:** Robert Grgac, Jan Rozsypal, Lauren Des Marteaux, Tomáš Štětina, Vladimír Koštál

## Abstract

Ability to survive freezing of extracellular body fluids evolved in several species of vertebrate ectotherms, many plants, and occurs relatively often in freeze-tolerant insects. Most of the multicellular organisms, however, are freeze-sensitive. Here we test coupled hypotheses postulating that: (i) irreversible denaturation of proteins and loss of integrity of biological membranes are two ultimate molecular mechanisms of freezing injury in freeze-sensitive insects; and (ii) seasonally accumulated small cryoprotective molecules (CPs) protect the proteins and membranes against the injury in freeze-tolerant insects. We show that seven different enzymes exhibit no or only partial loss of activity upon lethal freezing stress applied *in vivo* to whole freeze-sensitive larva of drosophilid fly, *Chymomyza costata*. In contrast, the enzymes lost activity when extracted and frozen *in vitro* in a diluted buffer solution. This loss of activity was fully prevented by adding to buffer relatively low concentrations of a wide array of different compounds including *C. costata* native CPs, other metabolites, bovine serum albumin (BSA), and even biologically inert artificial compounds Histodenz and Ficoll. Next, we show that the plasma membranes of fat body cells lose integrity when frozen *in vivo* in freeze-sensitive but not in freeze-tolerant larvae. Freezing fat body cells *in vitro*, however, resulted in loss of membrane integrity in both freeze-sensitive and freeze-tolerant larvae. Different additives showed widely different capacities (from none to high) to protect membrane integrity when added to *in vitro* freezing medium. A complete rescue of membrane integrity was observed for a mixture of proline, trehalose and BSA.

**Significance statement:** Here we suggest that insect soluble enzymes are not primary targets of freezing injury. They are not inactivated in freeze-sensitive insects exposed to lethal freezing stress as they are sufficiently protected from loss of activity by complex composition of native biological solutions. Next we show that cell plasma membranes are likely targets of freezing injury. The membranes lose integrity in freeze-sensitive insects exposed to freezing stress, while their integrity is protected by accumulated small cryoprotective molecules, and also by proteins, in freeze-tolerant insects.

## Introduction

Freezing of body fluids is lethal for most multicellular organisms though many plants, invertebrates and also a few amphibians and reptiles evolved capacity to survive internal ice formation - they are freeze tolerant (1-3). The freeze tolerance is particularly widespread among insects (4). All freeze tolerant organisms rely on extracellular ice formation while formation of ice inside their cells is (with exceptions) lethal (5). Extracellular ice causes osmotic outflow of water from cells, which results in cell shrinkage and concentration of cytosolic solutions (6). Consequently, the freeze-dehydrated cells are exposed to a combination of mechanical stress, low temperatures, decreasing activity of liquid water, and increasing levels of protons, metal ions or other chemical perturbants, which all together threaten conformational stability of proteins, lipid bilayers and other macromolecular complexes (7).

Widely-accepted consensus exists among environmental physiologists and low temperature biologists that the irreversible denaturation of proteins and loss of integrity of biological membranes are two important mechanisms of lethal freezing injury to various organisms (6, 8-11). Another almost unison agreement exists over the role of low molecular mass cryoprotective molecules (CPs), such as sugars, polyols, amino acids and their derivatives, that are accumulated by naturally freeze-tolerant organisms: the CPs are believed to protect proteins and membranes against freezing injury (1, 12-15). Accordingly, also in insect cold hardiness literature, the proteins and membranes are often mentioned simultaneously as two most likely targets of cold and freezing injury and their stabilization by seasonally accumulated CPs is accepted as a highly plausible hypothesis (1, 16-19).

The CPs are believed to help primarily by affecting the *colligative* properties of biological solutions - specifically by increasing osmolarity and thereby reducing the amount of ice formed at any given subzero temperature which, in turn, alleviates all mechanical, freeze-dehydration, and solute concentration stresses (6, 20-23). In addition, various *non-colligative* mechanisms of cryoprotection were hypothetically ascribed to insect CPs (1, 16, 23), and were also supported by indirect experimental evidence (24-26). The non-colligative mechanisms are theoretically based on noncovalent interactions between water molecules, de-stabilizing solutes (perturbants), stabilizing solutes (CPs), and stabilized proteins and lipid bilayers (reviewed in: (27-29). The protective effects of CPs were empirically demonstrated to occur in numerous *in vitro* experiments when isolated proteins or lipid bilayers were exposed to freezing or other stresses (27, 30-38). It has always been a tough challenge, however, to demonstrate that these *in vitro*-observed stabilizing effects of CPs do participate in cryoprotection *in vivo* – during the extracellular freezing of a freeze-tolerant organism. Here we aim to fill this gap in knowledge using the drosophilid fly, *Chymomyza costata* as a model.

The larvae of *C. costata* have two distinct seasonal phenotypes: active, warm acclimated larvae in summer, and diapause, cold acclimated larvae in winter (39). While active larvae are relatively freeze sensitive, the diapause larvae belong to the most cold-hardy animals known, can survive freezing of all osmotically active water in their body (68% water turned to ice crystals) down to -75°C, and survive even after long-term (18 mo) cryopreservation in liquid nitrogen (25, 40-43). In our previous work (26), we identified potential components of the native mixture of CPs that seasonally accumulate in hemolymph of diapause larvae (proline, trehalose, glutamine, asparagine, and betaine, to name the five most abundant molecules). We described the colligative effects of accumulated CPs on temperature-driven dynamism of water/ice transition (25), and demonstrated the existence of synergies between effects of different CPs (26).

Here we test the hypothesis that the seasonally accumulated CPs act as non-colligative stabilizers of soluble enzymes and plasma membranes of *C. costata* during freezing stress. We show that the soluble enzymes residing in their native biological solutions are well protected against freezing injury even in the freeze-sensitive phenotype of larvae, and, consequently, do not need any additional protection by seasonally accumulated CPs. In contrast, we found that the fat body cell plasma membrane is highly sensitive to freezing injury in both larval phenotypes, but its integrity during freezing stress is protected by CPs, specifically by proline and trehalose, and also by proteins that seasonally accumulate in hemolymph of the freeze-tolerant phenotype.

## Results

### Freezing stress kills freeze-sensitive larvae but does not inactivate their soluble enzymes

In the first series of experiments, we killed the larvae of *C. costata* of a freeze-sensitive phenotype by exposing them to SLOW inoculative freezing to -30°C (for schematic outline of experiment, see Fig. S1a). Immediately upon thawing, we assessed the post-freezing activities of select soluble enzymes extracted from different tissues. In these experiments, enzymes were frozen *in vivo* and unfrozen larvae of the same phenotype served as controls.

Figure 1 shows the examples of three different enzymes (glucose 6-P dehydrogenase, G6PDH; citrate synthase, CS; and lactate dehydrogenase, LDH). Four additional enzymes are shown in Fig. S2. Lethal freezing stress *never* caused complete loss of enzyme activity. Statistically significant decreases of activity were observed only in two cases: G6PDH in muscle (Fig 1a, decrease of activity by 32%) and amylases and maltases in midgut (Fig. S2c, decrease by 28%). In contrast, the activity of prophenoloxidase in hemolymph was considerably stimulated by freezing stress (Fig. S2a).

**Fig. 1:**
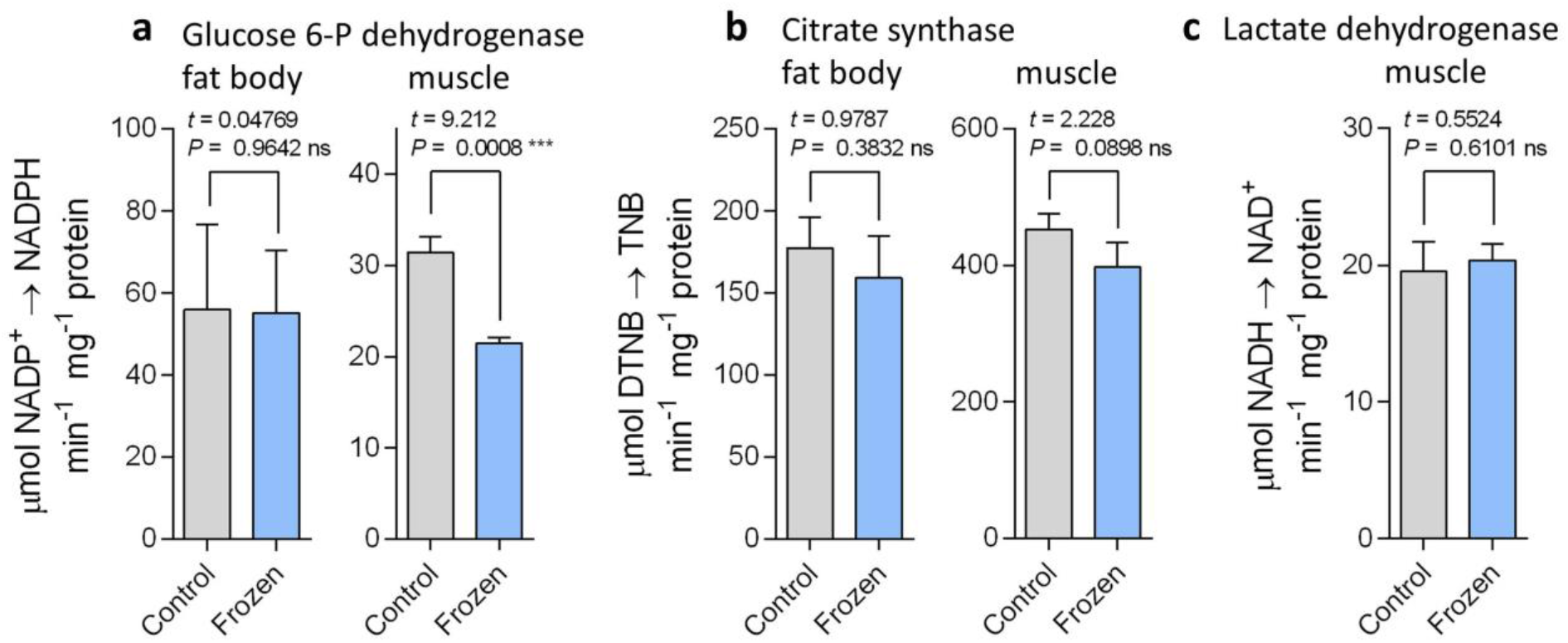
The enzyme activities persist after organismally-lethal freezing stress. The larvae of freeze-sensitive phenotype of *Chymomyza costata* were killed by freezing to -30°C (‘Frozen’). The ‘Control’ larvae were not frozen. The activities of three different enzymes: (**a**) glucose 6-phosphate dehydrogenase; (**b**) citrate synthase and; (**c**) lactate dehydrogenase were measured in total protein extracts of the fat body and muscle tissues dissected from Control and Frozen larvae right upon melting. Each column shows the mean + S.D. of the enzyme activity (*n* = 3, each replicate represents a pool of 30 larval tissues). The enzyme activities in Control vs Frozen larvae were compared using unpaired, two-tailed *t* test (*t* statistics and *P* values are shown: ns, not significant; *** significant difference; GraphPad Prism v. 6.07). For examples of other enzymes, see Figs. S2, S3.

In order to exclude that *C. costata* is exceptional in its ability to *in vivo* stabilize its soluble enzymes, we subjected two other insects to the same slow inoculative freezing as applied to larvae of *C. costata*: the larvae of the vinegar fly, *Drosophila melanogaster*; and the tibial levator muscle dissected from femur of the hind leg of the adult locust, *Locusta migratoria*. Immediately after thawing the specimens, we extracted total proteins from muscle tissue and assessed the activity of LDH: it was by 41% higher in the muscle of frozen than in control larvae of *Drosophila* (Fig. S3a), while no loss of LDH activity was observed in frozen muscle of locust (Fig. S3b).

### Freezing stress inactivates enzymes *in vitro*

In the second series of experiments, we exposed to freezing stress the total protein extracts of the fat body or muscle tissue of the freeze-sensitive larvae of *C. costata*. In these experiments, the enzymes experienced freezing stress in a diluted aqueous solution of 20 mM imidazole buffer (i.e. *in vitro*). The protein extracts were divided to aliquots: one aliquot serving as control (no additive, unfrozen), the other aliquots were exposed to freezing stress in presence of different concentrations of different additives. We used two different freezing protocols: (i) FAST, which simulates routine laboratory practice when a protein extract in Eppendorf tube is simply moved to a deep freezer set to -75°C; and (ii) SLOW, which simulates ecologically relevant, slow inoculative freezing (to compare FAST and SLOW *in vitro* freezing protocols, see Fig S1b). In both FAST and SLOW protocols, we invariably observed complete (or almost complete) loss of activity in three different enzymes, G6PDH, CS, and LDH. We are not presenting these results in an independent figure as this would only show a difference between 100% enzymatic activity in unfrozen control vs. 0% (or close to 0%) activity in frozen extract. Instead, these results are presented as initial points (concentration of additive = 0 on *x* axis) in Figs. 2 and 3 and in related supplementary figures in SI file.

**Fig. 2.**
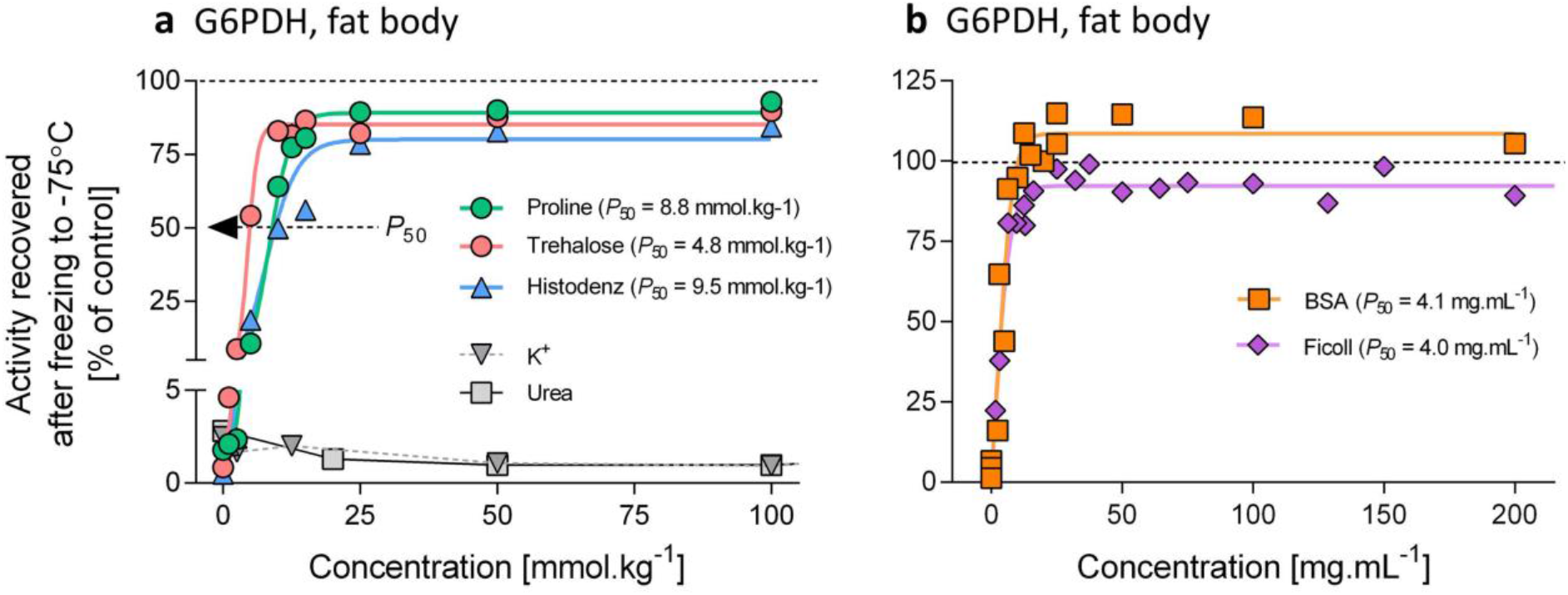
The enzyme activity is protected by low concentrations of various additives to the *in vitro* freezing solution. The total proteins were extracted from the fat body tissue of freeze-sensitive phenotype of *Chymomyza costata* larvae and the activity of glucose 6-phosphate dehydrogenase (G6PDH) were measured prior to (control) and after (test) FAST freezing to -75°C (see Fig. S1b for protocol). The test aliquots were augmented with different additives: (**a**) proline, trehalose, Histodenz, K^+^, and urea; (**b**) BSA and Ficoll; administered at different concentrations (see *x* axes). The activity in the unfrozen aliquot served as control = 100%. Freezing without additive (concentration = 0) caused complete loss of enzyme activity. Different additives protected the enzyme from loss of activity upon freezing at relatively low concentrations (*P*_50_ is a cryoprotective concentration that allows recovering 50% of control, pre-freezing activity). The K^+^ and urea showed no cryoprotective effects. For other additives and enzymes, see Fig. S5.

**Fig. 3.**
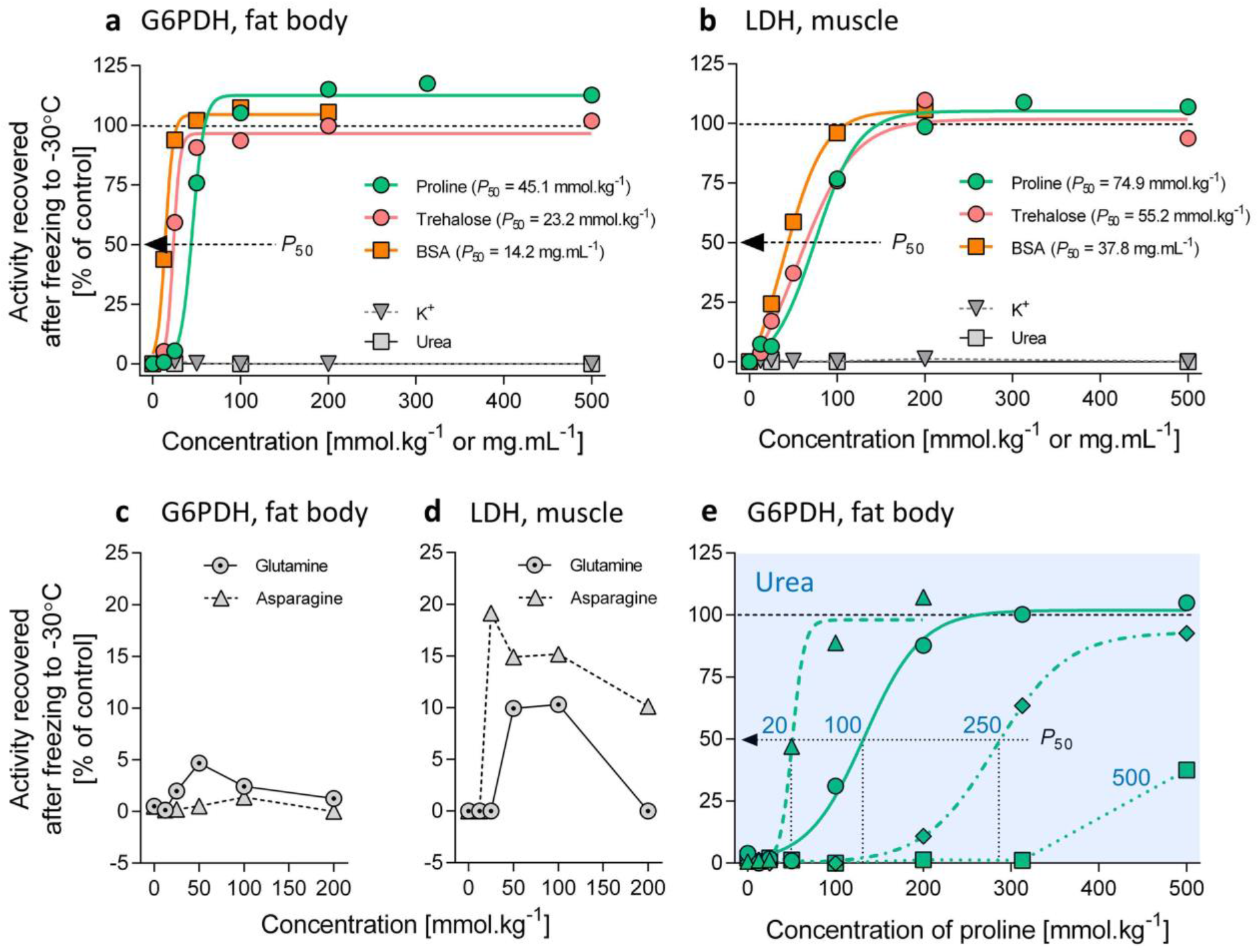
Slow rate of *in vitro* freezing in the absence and presence of chemical perturbants. The total proteins were extracted from the fat body or muscle tissues of freeze-sensitive phenotype of *Chymomyza costata* larvae and the activities of glucose 6-phosphate dehydrogenase (G6PDH) and lactate dehydrogenase (LDH) were measured prior to (control) and after (test) SLOW inoculative freezing to -30°C (see Fig. S1b for protocol). The test aliquots were augmented with different additives administered at different concentrations (see *x* axes). The activity in the unfrozen aliquot served as control = 100%. Freezing without additive (concentration = 0) caused complete loss of enzyme activity. Different additives protected the enzyme from loss of activity upon freezing at relatively low concentrations (*P*_50_ is a cryoprotective concentration that allows recovering 50% of control, pre-freezing activity). (**a, b**) Proline, trehalose, and BSA protected the enzymes from loss of activity at relatively low *P*_50_ concentrations. The K^+^ and urea showed no cryoprotective effects. For other additives, see Fig. S6a. (**c, d**) Glutamine and asparagine showed no or only weak ability to protect the enzymes. (**e**) Simultaneous presence of urea (perturbant) and proline (CP) during SLOW freezing resulted in shifting the *P*_50_ values for proline toward higher concentrations (for other combinations of CPs and chemical perturbants, see Fig. S6b-d).

### Low concentrations of CPs are sufficient to protect the activity of enzymes frozen *in vitro*

Next, we applied different additives to 20 mM imidazole buffer in order to assess whether this change in composition of the *in vitro* freezing solution can protect the enzymes of *C. costata* from loss of activity upon freezing stress. We searched for the additive concentration (*P*_50_) that ensures 50% recovery of initial (unfrozen control) enzyme activity after freezing stress. In preliminary experiments, we verified that the incubation of protein extract at 0°C (without freezing) for 1h, with or without CP additives, have no or little effect on G6PDH activity (Fig. S4a, b). The incubation with potential chemical perturbants was also tested. The loss of G6PDH activity was seen only with urea administered at concentrations higher than 500 mmol.kg^−1^ (Fig. S4c).

We found that very low concentrations of different additives were sufficient to protect G6PDH extracted from fact body cells from loss of activity upon FAST *in vitro* freezing (Fig. 2; Fig. S5). Five components of *C. costata* native cryoprotectant mixture showed *P*_50_ values ranging between 4.8 mmol.kg^−1^ (trehalose, Fig. 2a) and 13.9 mmol.kg^−1^ (asparagine, Fig. S5a). Biologically inert low-molecular-weight additive Histodenz (*P*_50_, 9.5 mmol.kg^−1^, Fig. 2a) and general-use cryoprotectant glycerol (*P*_50_, 13.1 mmol.kg^−1^, Fig. S5a) showed similar cryoprotective abilities. The macromolecules BSA and Ficoll (biologically inert additive) also protected G6PDH at very low concentrations: BSA, *P*_50_, 4.1 mg.mL^−1^; and Ficoll, 4.0 mg.mL^−1^ (Fig. 2b). In contrast, urea and K^+^ failed to protect G6PDH at any concentration assessed (Fig. 2a). Despite being perfect cryoprotectants, neither proline nor Ficoll, were able to protect G6PDH from almost complete loss of activity upon heat stress (Fig. S5b).

We conducted similar experiments (though less extensive) as described above for two other enzymes with basically similar results: CS extracted from fat body and LDH extracted from muscle of *C. costata* larvae lost their activities completely upon FAST *in vitro* freezing stress without additives. However, very low concentrations of proline (*P*_50_, 7.2 and 8.0 mmol.kg^−1^), trehalose (*P*_50_, 7.3 and 4.6 mmol.kg^−1^), or BSA (*P*_50_, 4.9 and 1.5 mg.mL^−1^) were sufficient to protect the CS and LDH, respectively, from loss of activity upon FAST *in vitro* freezing (Fig. S5c, d).

### Slow rate of *in vitro* freezing in the presence of chemical perturbants

Next, we exposed the G6PDH and LDH enzymes *in vitro* to a more ecologically-relevant, SLOW inoculative freezing protocol (Fig S1b). The *P*_50_ concentrations for all protective additives (proline, trehalose, betaine, glycerol, Histodenz, BSA, and Ficoll) were invariably higher in the SLOW than in the FAST protocol. For instance, 45.1 mmol.kg^−1^ of proline was needed to protect 50% G6PDH activity in the SLOW protocol, while 8.8 mmol.kg^−1^ was sufficient in the FAST protocol. The urea and K^+^ failed to protect the G6PDH from loss of activity (Fig. 3a, b; Fig S6a). Furthermore, glutamine and asparagine, though behaving as perfect cryoprotectants in the FAST protocol, showed only negligible (G6PDH, Fig. 3c) or weak (LDH, Fig. 3d) ability to protect the enzymes in the SLOW protocol.

Further, we assessed the capacity of proline and trehalose to stabilize G6PDH during SLOW inoculative freezing stress in the presence of chemical perturbants of the enzyme activity such as urea and K^+^ (glutamine was also tested as potential perturbant as it failed to protect the enzyme). When urea (> 100 mmol.kg^−1^) was present in the *in vitro* freezing medium, considerably higher *P*_50_ concentrations of proline (Fig. 3e) or trehalose (Fig. S6b) were needed to protect the enzyme from loss of activity than in the absence of urea. Presence of K^+^ or glutamine had no effect on *P*_50_ concentrations of proline (Fig. S6c, d). We used differential scanning calorimetry to measure the amount of ice fraction, which allowed us to estimate the increase in concentration of urea caused by freezing the osmotically active water in solutions combining urea plus CPs (proline or trehalose) (Fig. S7a). This analysis showed that the cryoprotective effect of CPs in the presence of urea can be explained partially by decreasing the freeze-concentration of urea and partially by counteracting the chemical perturbance caused by high concentrations of urea (see Fig. S7 for more details).

### Unprotected plasma membrane loses integrity upon freezing stress both *in vivo* and *in vitro*

Using Trypane Blue assay, we assessed the integrity of the plasma membranes of *C. costata* larval fat body cells exposed to SLOW inoculative freezing stress either *in vivo* (whole larvae frozen) or *in vitro* (the dissected fat body tissue exposed to freezing in Schneider’s *Drosophila* medium) (for schematic outline of experiment, see Fig. S1c). The results for active (freeze-sensitive) vs diapause (freeze-tolerant) larvae were different. In active larvae, 64% of fat body cells stained blue (indicating that the plasma membrane lost integrity) after freezing *in vivo*, and 94% stained blue after freezing *in vitro* (Fig. 4a). In diapause larvae, only 2.6% of fat body cells cells stained blue after freezing *in vivo*, while 95% stained blue after freezing *in vitro* (Fig. 4b). These results suggest that the hemolymph of diapause larvae contains components *necessary and sufficient* for protection of the diapause cell plasma membrane integrity upon freezing stress.

**Fig. 4:**
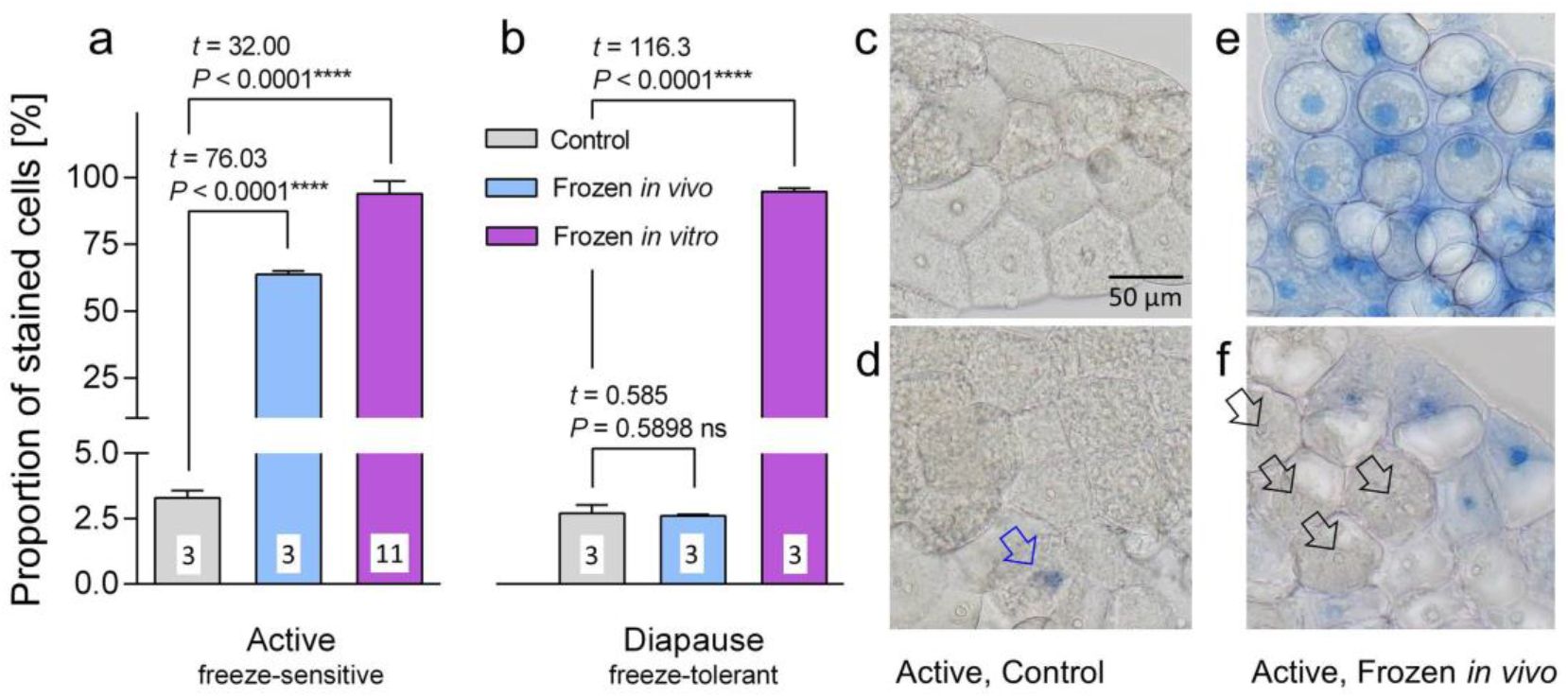
The integrity of unprotected fat body cell plasma membrane is compromised by freezing stress. **(a**) Active, freeze-sensitive and (**b**) diapause, freeze-tolerant phenotypes of *Chymomyza costata* larvae were used for experiments on maintenance of membrane integrity upon SLOW inoculative freezing to -30°C. Unfrozen larvae (grey columns) served as controls. Either whole larvae were exposed to freezing stress (*in vivo*, blue columns; see Fig. S1a for protocol) or the dissected fat body tissues were frozen in the Schneider’s *Drosophila* medium (*in vitro*, violet columns, see Fig. S1c for protocol). After melting, the integrity of fat body cell plasma membrane was assessed using Trypane Blue assay (the cells with compromised membrane integrity stain blue). Each column shows the mean + S.D. of blue staining (*n* = 3 or 11, each replicate represents a pool of 10 larval tissues). The differences in staining proportion were compared using unpaired, two-tailed *t* tests (*t* statistics and *P* values are shown: ns, not significant; **** significant difference; GraphPad Prism v. 6.07). The examples of staining patterns in active, freeze-sensitive larvae are shown in (**c**) control, unfrozen larva, no staining visible; (**d**) other control, unfrozen larva, one cell nucleus stained (blue arrow); (**e**) *in vivo* frozen larva, all cells stained; (**f**) other *in vivo* frozen larva, most cells stained but there is an ‘island’ of unstained cells (white arrows). The micrographs were taken using Olympus SZX12 binocular microscope; the 50 µm bar shown in (c) applies to all four micrographs.

### Proline, trehalose and BSA protect membrane integrity upon freezing stress *in vitro*

Next, we assessed whether different additives to the Schneider’s medium can protect the plasma membranes of fat body cells of the freeze-sensitive larval phenotype from loss of integrity upon freezing stress applied to dissected tissue *in vitro*. Prior to this experiment, we verified that the additives themselves (without freezing stress) do not compromise the membrane integrity (Fig. S8a). We found that only two of five tested components of the *C. costata* native cryoprotectant mixture were partially effective in membrane cryoprotection. Proline decreased the ratio of blue-stained cells from 94% (control) to 78%, 74% and 85% when applied at concentrations of 313, 500 and 750 mmol.kg^−1^, respectively; and trehalose decreased the ratio of blue-stained cells to 85% and 78% when applied at concentrations of 250 and 500 mmol.kg^−1^ (Fig. 5a). Other three components of native cryoprotectant mixture of *C. costata* (glutamine, asparagine, and betaine) were ineffective in membrane protection (Fig. S8c) as were also saccharose, Histodenz, and Na^+^ (Fig. 5a). Glycerol, which is not natively accumulated by *C. costata* larvae, was even more effective in membrane cryoprotection than proline and trehalose (Fig. S8b). The macromolecular additive BSA showed cryoprotective effect at concentrations ranging from 25 to 200 mg.mL^−1^, while Ficoll was marginally effective only at the highest concentration applied (200 mg.mL^−1^, ANOVA *P* = 0.0477) (Fig. 5b). The cryoprotective effects of different CPs applied at the same osmotic concentration (500 mOsmol.kg^−1^) drastically differed (Fig. S8d), which demonstrates that the effect is not explainable simply based on colligative properties of the augmented-Schneider’s medium. Examples of Trypane blue staining of *in vitro* frozen fat body cells are shown in Fig. S9.

**Fig. 5.**
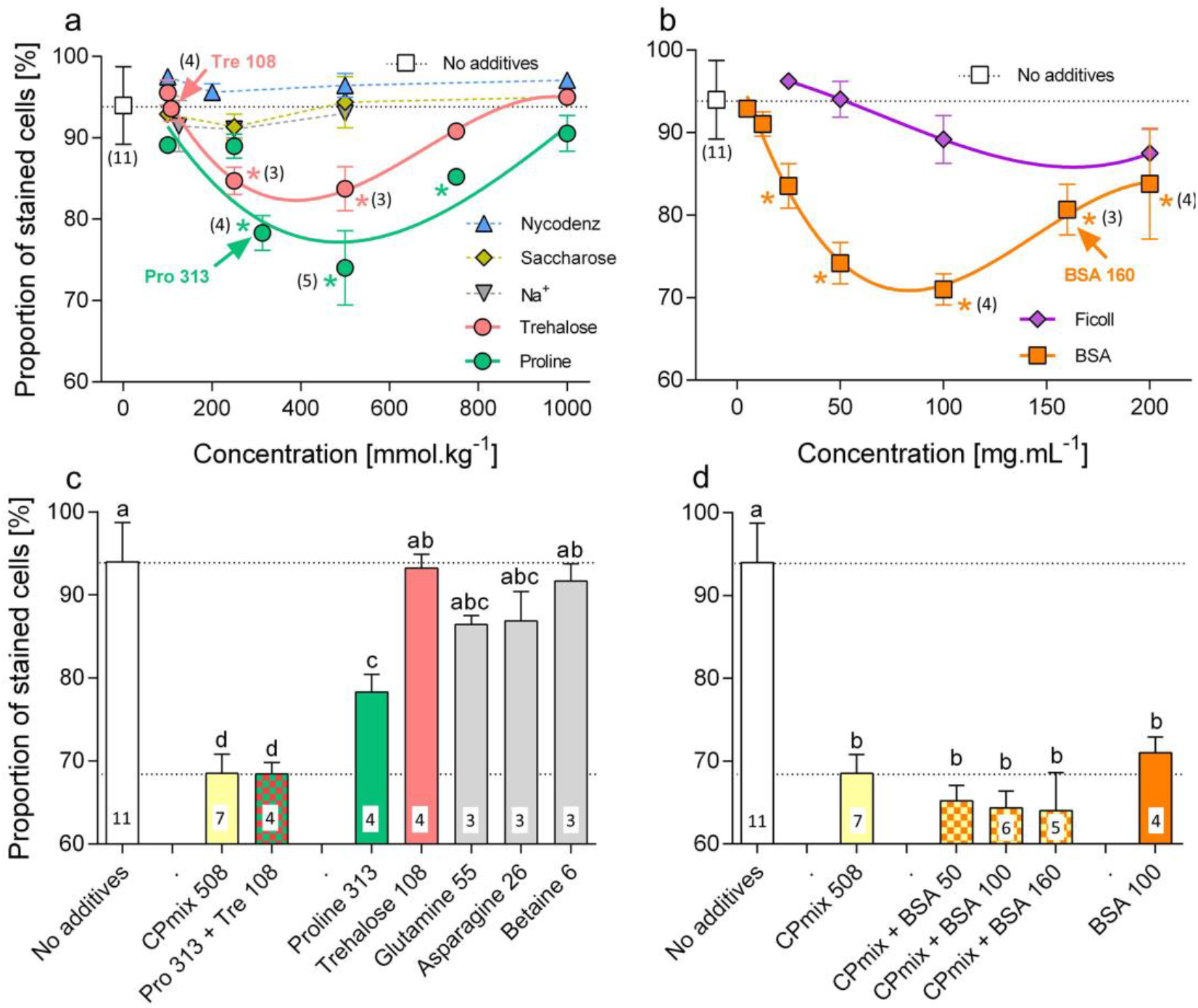
The membrane integrity upon freezing stress is partially protected by high concentrations of proline, trehalose, and BSA. The dissected fat body tissues of freeze-sensitive phenotype larvae of *Chymomyza costata* were exposed to freezing stress *in vit*ro, in Schneider’s solution augmented by different additives at different concentrations. The control tissues (empty squares in (a, b) or empty columns in (c, d) were exposed to freezing stress without additive. After melting, the integrity of fat body cell plasma membrane was assessed using Trypan Blue assay (see *y* axes). The panels (**a, b**) compare the cryoprotective effects of different concentrations (see *x* axes) of different additives (for other additives, see Fig. S8b, c). The concentrations corresponding to physiological levels observed in the hemolymph of freeze-tolerant larvae are indicated by arrows flanked by numbers (the concentrations).; the third order polynomial curves or dashed lines are used to visually connect different concentrations of an additive. The panels (**c, d**) compare the cryoprotective effects of different additives added singly or in various mixtures (for other combinations, see Fig. S8d, e). The concentrations are indicated (313, 108, 55, 26, and 6 mmol.kg^−1^ for CPs; 50, 100, 160 mg.mL^−1^ for BSA; 508 mmol.kg^−1^ for CPmix of five CPs). Each point or column shows the mean ± S.D. of blue staining in at least two replicates, while treatments replicated three or more times are indicated by numbers (*n* = 3-11, each replicate represents mean staining in a pool of 10 larval tissues). The differences in staining proportions were compared using one-way ANOVA tests followed by Bonnferoni’s multiple comparison tests (GraphPad Prism v. 6.07). In (a, b), the additive-augmented-media were compared to control) and statistically significant differences are flanked by *; in (c, d) each column was compared with every other column and the columns flanked by different letters are statistically different.

Looking at interactions between CPs, we compared cryoprotective effects of different CPs applied singly with those of various mixtures. We prepared artificial mixture of five putative CPs (CPmix 508) simulating the physiological concentrations of individual components as they were detected in hemolymph of freeze-tolerant phenotype larvae of *C. costata* (i.e. proline, 313 mmol.kg^−1^; trehalose, 108 mmol.kg^−1^; glutamine, 55 mmol.kg^−1^; asparagine, 26 mmol.kg^−1^, and betaine, 6 mmol.kg^−1^; in total 508 mmol.kg^−1^). After freezing in the CPmix 508-augmented Schneider’s medium, the ratio of blue-stained cells decreased to 68.5 % (Fig. 5c). The same cryoprotective effect (68.5 %), however, was observed for a mixture combining just two major CPs, proline (313 mmol.kg^−1^) and trehalose (108 mmol.kg^−1^). Comparing these results to the cryoprotective effects of different CPs applied singly, we can conclude that: (i) proline was the only component that significantly decreased the ratio of blue-stained cells when applied at its physiological concentration corresponding to freeze-tolerant hemolymph; (ii) there was a synergism between proline and trehalose in their cryoprotective effects (simple addition of the individual effects of two components does not explain the cryoprotective effect observed in their mixture) (Fig. 5c). We further confirmed the synergism between proline and trehalose in the experiment where individual components were removed from the CPmix 508 one by one. Removing glutamine; asparagine, or betaine had practically no impact on cryoprotective effect of the reduced CPmix. Removing proline or trehalose, however, significantly decreased the cryoprotective effect of the reduced CP mix (Fig. S8e). (iii) Combining the CPmix 508 with BSA at different concentrations, no synergy was achieved, but a weak, statistically insignificant additive effect was observed (Fig. 5d).

### Complete rescue of membrane integrity by CPs and BSA in the freeze-tolerant phenotype larvae

The plasma membranes of fat body cells of the freeze-tolerant larval phenotype also lose integrity when the cells are frozen in the plain Schneider’s medium *in vitro* (Figs. 4b, S8f). The CPs proline or trehalose, when added singly to Schneider’s medium at their respective physiological concentrations, reduced the Trypan Blue staining significantly from 95% to 66% or 73%, respectively. An additive effect, or a weak synergism, was observed for a mixture of proline and trehalose (reduction of staining to 31%), which was practically the same cryoprotective effect as observed for complete CPmix 508 (reduction of staining to 33%). The BSA appeared as highly potent cryoprotective agent as it reduced the Trypan Blue staining to 10% when applied at concentration corresponding to physiological concentration of total hemolymph proteins in freeze-tolerant larvae. We achieved a complete rescue of plasma membrane integrity upon freezing stress (reduction of staining below 3%) when a mixture of just three components, proline, trehalose, and BSA, was added to Schneider’s medium (Fig. S8f).

## Discussion

Here we provide an experimental test of the widely accepted hypothesis that sees the irreversible protein denaturation and loss of integrity of biological membranes as two most important molecular mechanisms of freezing injury in freeze-sensitive insects, while the freeze tolerant-insects are hypothesized to prevent this injury using accumulated cryoprotective solutes (1, 16-18) – for exact quotations, see SI Supplementary Discussion.

### Insect soluble enzymes *in vivo* are not the primary targets of freezing injury and do not need protection by accumulated CPs

To our best knowledge, the insect cold hardiness literature describes only single direct experiment on insect enzyme activity measured prior to and after freezing stress – the citrate synthase in the fat body cells of lethally frozen *C. costata* larvae was not inactivated (44). This isolated observation prompted us to extend the assays on *in vivo* freeze-inactivation/stability to other insect enzymes. Here, we show that seven randomly chosen soluble enzymes representing different biological solutions (hemolymph, alimentary canal, extracellular matrix, cytosol and mitochondrial matrix of muscle and fat body cells) of three different freeze-sensitive insects (i.e. insects not protected by any accumulated CPs) were not inactivated during organismally-lethal freezing stress. Considering the possible cause of discrepancy between hypothesis and empirical results, we think that the high sensitivity of enzymes exposed to freezing stress *in vitro* might the major source of misinterpretation.

### Soluble enzymes are prone to loss of activity upon freezing *in vitro* in a diluted aqueous solution

We observed complete loss of activities in *C. costata*’s G6PDH, LDH and CS when they were frozen *in vitro*, in tissue extracts diluted in 20 mM imidazole buffer. This observation is in agreement with relatively rich literature on vertebrate (30, 36, 45) and insect enzymes (35). Our experiments show that the loss of enzyme activity is preventable by adding relatively low concentrations of various micro-solutes (including biologically inert Histodenz) into the freezing medium. This is in agreement to earlier study by Storey et al. (35) who found almost exactly same low *P*_50_ values (7 – 25 mM) for a range of sugars, polyols and amino acids protecting the G6PDH from freezing inactivation in two other insects, *Eurosta solidaginis* and *Epiblema scudderiana* and also yeast. Such low concentrations of various micro-solutes, however, occur in almost every organism and/or cell type. Even some vertebrate soluble enzymes were observed to survive the freeze-thaw cycle when frozen *in situ*, inside the tissue: e.g. succinate dehydrogenase and cytochrome oxidase in rat heart (46); or sarcoplasmic ATPase and aldolase in fish meat, which lost activity significantly only upon prolonged frozen-storage (47, 48). Higher concentrations of CPs (in a range of hundreds mM) were needed to protect the purified mammalian enzymes (G6PDH, LDH and other commercially available enzymes) than insect enzymes from loss of activity upon *in vitro* freezing ((30, 45) and elder literature cited therein). Even the mammalian enzymes, however, were well protectable by relatively low concentrations (in a range of 0.02 – 0.25 mg.mL^−1^) of macromolecular compounds such as polyethylene glycol (PEG), BSA or by increasing the concentration of the enzyme itself (self-protection) (30, 45). Accordingly, here we also observed that BSA or Ficoll at concentrations as low as 4 – 40 mg.mL^−1^ were sufficient to perfectly protect the insect enzymes from loss of activity upon freezing. Typical protein concentrations in biological solutions, however, are higher by one or two orders (in a range of 50 – 400 mg.mL^−1^) (49).

Collectively, soluble enzymes seem to be sufficiently protected against loss of activity when frozen in their native biochemical environment, in a biological solution which is ‘crowded’ by various micro-solutes and macromolecules (50, 51). Of course, this statement is open to scientific falsification as we can hardly exclude that some other enzymes, not included in this study, are more sensitive to freezing stress. Moreover, there are several reasons for cautiousness when extending the validity of our hypothesis from *soluble enzymes* to *all proteins*. We corroborate these reasons in the SI section: ‘Do proteins in general need stabilization by CPs during freezing stress?’

### Cell plasma membrane needs protection by CPs and proteins in order to maintain integrity during freezing stress

In agreement with widely accepted hypothesis on molecular mechanisms of freezing injury, here we confirm experimentally that the cell plasma membranes of the freeze-sensitive phenotype larvae of *C. costata* lose their barrier function upon *in vivo* lethal freezing stress while almost no loss of membrane integrity is observed in the freeze-sensitive phenotype. The plasma membrane is considered as primary target of cold and freezing injury also in plants (11) and mammalian cells (6, 52).

One salient observation of this work points to the hemolymph of freeze-tolerant phenotype of *C. costata* larva, which apparently contains components both *necessary and sufficient* for protection of the plasma membrane integrity upon freezing stress. In earlier study (26), we identified the micro-molecular components of the freeze-tolerant phenotype hemolymph and, in this study, we tested their cryoprotective abilities in combination with macromolecular BSA. A combination of three additives to Schneider’s medium, i.e. proline, trehalose, and BSA, appeared as sufficient to completely rescue the membrane integrity upon freezing stress applied *in vitro* to fat body cells of freeze-tolerant larvae. The same mixture also significantly reduced the injury caused by freezing stress to membranes of fat body cells of the freeze-sensitive phenotype. Of the three additives, the BSA was the most effective in membrane cryoprotection for both freeze-sensitive and freeze-tolerant phenotypes. We used BSA in concentration that corresponds to the physiological concentration of total proteins in the hemolymph of *C. costata* larvae (i.e. 100 and 160 mg.mL^−1^ for freeze-sensitive and -tolerant phenotypes, respectively). Drosophilid larvae accumulate serum proteins late in their ontogeny (prior to metamorphosis) (53) and many insects, including larvae of *C. costata*, do accumulate even more of them prior to diapause, as a source of amino acids for post-diapause development (54, 55). The BSA and other membrane-non-permeable compounds such as saccharose, Ficoll, PEG, or hydroxyethyl starch are often added to cryopreservation solutions for mammalian sperm (56) where they likely help to maintain membrane integrity indirectly via affecting the thermal transitions in extracellular medium including kinetics of growth and morphology of ice crystals and glass transition temperatures (57, 58), and/or by reducing the membrane lipid peroxidation by reactive oxygen species (59). The exact mechanism behind high effectiveness of BSA as *C. costata*’s membrane integrity protectant, however, remains currently unknown. Other two cell membrane non-permeable compounds (saccharose and Ficoll) showed no or negligible cryoprotective effects toward *C. costata* fat body cell membranes. Additional research is required to determine whether different CPs permeate plasma membrane and how this impacts its integrity in the frozen state.

Our functional assays further corroborate results of earlier studies where insect fat body cells were exposed to freezing *in vitro* and the integrity of their plasma membrane was then checked using vital dyes: (i) fat body cells of the larvae of *Eurosta solidaginis* were frozen to -25°C in Grace’s insect medium and the augmentation of Grace’s medium with 1M glycerol increased the proportion of cells with intact plasma membrane from less than 20% to 80% (60); (ii) fat body cells of the cricket *Gryllus veletis* were frozen to -12°C (warm-acclimated crickets) or -16°C (cold-acclimated crickets) in Grace’s medium - plain or augmented with different CPs: the augmentations with myo-inositol, trehalose, and glycerol significantly increased the proportions of cells that survived freezing with intact plasma membrane (24). The later study also suggested that individual CPs differentially impact survival in the frozen state, are not interchangeable, and likely function non-colligatively in insect freeze tolerance. Similarly as in the present study, no CP or combination was sufficient to confer high freeze tolerance to cells of warm-acclimated crickets, whereas in cold-acclimated crickets the proportion of cells with intact membrane after freezing stress increased to approximately 75% after adding the combination of myo-inositol plus trehalose or glycerol alone (24).

## Conclusions

Our results suggest that the seasonal accumulation of small cryoprotective molecules - a characteristic physiological response to cold-acclimation observed in many insect species - was evolutionarily driven, at least partially, by the necessity to protect plasma cell membranes against the loss of integrity during cold and freezing stress. In addition, membrane integrity seems to be importantly supported by accumulated hemolymph proteins. In contrast, our results do not support the hypothesis that the stability of insect soluble enzymes would be endangered by cold and freezing stress. Though the enzymes are highly sensitive to freezing stress in diluted aqueous solutions *in vitro*, they show high stability when exposed to freezing *in vivo*, in their native biological solutions crowded by different solutes and macromolecules.

## Materials and Methods

For a detailed description of materials and methods, see SI Supplementary Methods.

### Insects

Two phenotypic variants of *C. costata* larvae were generated according to our earlier acclimation protocols (26): (i) active, warm-acclimated larvae; and (ii) diapause, cold-acclimated larvae. The active larvae have limited survival after freezing stress (none survives when frozen below -10°C), while practically all diapause larvae survive freezing down to - 75°C, and 42.5% survive and metamorphose into fit adults even after 18 months of cryopreservation in LN_2_ (25). Since we focused on the nature of damage caused by freezing stress, the freeze-sensitive, active larvae were used for most experiments. The diapause larvae were used for the experiments on integrity of plasma membrane upon freezing stress.

### General outline of experiments

The general rationale of all experiments was to expose either whole active, freeze-sensitive larvae (*in vivo*), or their dissected tissues (*in vitro*), or protein extracts from their tissues (*in vitro*), to lethal freezing stress of -30°C and, upon melting, assess: (i) whether the plasma membranes of their fat body cell lost integrity; (ii) whether their soluble enzymes lost activity; (iii) and whether these losses of activity/integrity can be prevented by different additives into the freezing medium. The general outline of all experiments is presented in Fig. S1.

### Enzyme activity and plasma cell membrane integrity assays

The activities of seven different enzymes were measured (as specified in SI Materials and Methods) prior to and after the freezing stress. Similarly, plasma membrane integrity prior to and after the freezing stress was assessed using Trypan Blue dye which stains interior of cells with compromised membrane barrier function. The *in vitro* experiments allowed us to manipulate the composition of freezing medium. For enzyme assays, 20 mM imidazole was the base of *in vitro* freezing medium, while Schneider’s *Drosophila* medium (Biosera, Nuaillé, France) was used for *in vitro* incubation and freezing of dissected tissues. The freezing media were augmented by additives in different concentrations. We searched for the additive concentration (*P*_50_) that ensures 50% recovery of initial (unfrozen control) enzyme activity. The additives included: (i) the components of native cryoprotective mixture accumulated in diapause, freeze-tolerant phenotype of *C. costata* larvae, i.e. proline, trehalose, glutamine, asparagine, and betaine (26); (ii) other additives widely used in cryobiology practice, i.e. glycerol, and saccharose; (iii) inorganic salts, NaCl and KCl; (iv) bovine serum albumin; (v) biologically inert compounds Histodenz and Ficoll. In addition, urea was used as chemical perturbant of native protein structure in some experiments (see Table S1 for complete list of additives).

## Acknowledgements

We thank Irena Vacková and Jaroslava Korbelová for maintenence of insect colonies. This study was supported by a Grantová Agentura České Republiky (GAČR) grant no. 19-13381S to VK.

## References

1. Storey KB & Storey JM (1988) Freeze tolerance in animals. Physiol. Rev. 68(1):27–84.

2. Asahina E (1969) Frost resistance in insects. Adv. Insect Physiol., (Elsevier), Vol 6, pp 1–49.

3. Pearce RS (2001) Plant freezing and damage. Ann. Bot. 87(4):417–424.

4. Sinclair BJ (1999) Insect cold tolerance: How many kinds of frozen? Eur. J. Entomol. 96:157–164.

5. Sinclair BJ & Renault D (2010) Intracellular ice formation in insects: unresolved after 50 years? Comp Biochem Physiol A 155(1):14–18.

6. Mazur P (1984) Freezing of living cells: mechanisms and implications. Am. J. Physiol. 247(3):C125–C142.

7. Muldrew K, Acker JP, Elliott JA, & McGann LE (2004) The water to ice transition: implications for living cells. Life in the frozen state, (CRC Press), pp 93-134.

8. Franks F & Hatley RH (1991) Stability of proteins at subzero temperatures: thermodynamics and some ecological consequences. Pure Appl. Chem. 63(10):1367–1380.

9. Dias CL, et al. (2010) The hydrophobic effect and its role in cold denaturation. Cryobiology 60(1):91–99.

10. Ramløv H (2000) Aspects of natural cold tolerance in ectothermic animals. Hum. Reprod. 15(5):26–46.

11. Steponkus PL (1984) Role of the plasma membrane in freezing injury and cold acclimation. Annu. Rev. Plant Physiol. 35(1):543–584.

12. Yancey PH & Siebenaller JF (2015) Co-evolution of proteins and solutions: protein adaptation versus cytoprotective micromolecules and their roles in marine organisms. J. Exp. Biol. 218(12):1880–1896.

13. Yancey PH (2005) Organic osmolytes as compatible, metabolic and counteracting cytoprotectants in high osmolarity and other stresses. J. Exp. Biol. 208(15):2819–2830.

14. Hochachka PW & Somero GN (2002) Biochemical adaptation: mechanism and process in physiological evolution (Oxford university press).

15. Somero G (1986) Protons, osmolytes, and fitness of internal milieu for protein function. Am. J. Physiol. Regul. Integr. Comp. Physiol. 251(2):R197–R213.

16. Lee REJ (2010) A primer on insect cold-tolerance. Low Temperature Biology of Insects, eds Denlinger DL & Lee REJ (Cambridge University Press).

17. Teets NM & Denlinger DL (2013) Physiological mechanisms of seasonal and rapid cold-hardening in insects. Physiol. Entomol. 38(2):105–116.

18. Toxopeus J & Sinclair BJ (2018) Mechanisms underlying insect freeze tolerance. Biol. Rev. 93:1891–1914.

19. Rozsypal J (2022) Cold and freezing injury in insects: An overview of molecular mechanisms. EJE 119(1):43–57.

20. Meryman H (1971) Cryoprotective agents. Cryobiology 8(2):173–183.

21. Lovelock J (1954) The protective action of neutral solutes against haemolysis by freezing and thawing. Biochem. J. 56(2):265–270.

22. Zachariassen KE (1985) Physiology of cold tolerance in insects. Physiol. Rev. 65(4):799–832.

23. Storey KB (1997) Organic solutes in freezing tolerance. Comp Biochem Physiol A 117(3):319–326.

24. Toxopeus J, Koštál V, & Sinclair BJ (2019) Evidence for non-colligative function of small cryoprotectants in a freeze-tolerant insect. Proc. R. Soc. B 286(1899):20190050.

25. Rozsypal J, Moos M, Šimek P, & Koštál V (2018) Thermal analysis of ice and glass transitions in insects that do and do not survive freezing. J. Exp. Biol. 221:170464.

26. Kučera L, et al. (2022) A mixture of innate cryoprotectants is key for freeze tolerance and cryopreservation of a drosophilid fly larva. J. Exp. Biol. 225(8):jeb243934.

27. Timasheff S (1992) A physicochemical basis for the selection of osmolytes by nature. Water and life, (Springer), pp 70–84.

28. Timasheff SN (2002) Protein-solvent preferential interactions, protein hydration, and the modulation of biochemical reactions by solvent components. Proceedings of the National Academy of Sciences 99(15):9721–9726.

29. Timasheff SN (1993) The control of protein stability and association by weak interactions with water: how do solvents affect these processes? Annu. Rev. Biophys. Biomol. Struct. 22(1):67–97.

30. Carpenter JF & Crowe JH (1988) The mechanism of cryoprotection of proteins by solutes. Cryobiology 25(3):244–255.

31. Arakawa T & Timasheff SN (1982) Stabilization of protein structure by sugars. Biochemistry 21(25):6536–6544.

32. Anchordoguy TJ, Rudolph AS, Carpenter JF, & Crowe JH (1987) Modes of interaction of cryoprotectants with membrane phospholipids during freezing. Cryobiology 24(4):324–331.

33. Crowe LM, Mouradian R, Crowe JH, Jackson SA, & Womersley C (1984) Effects of carbohydrates on membrane stability at low water activities. Biochim. Biophys. Acta (BBA)-Biomembr. 769(1):141–150.

34. Arakawa T & Timasheff S (1985) The stabilization of proteins by osmolytes. Biophys. J. 47(3):411–414.

35. Storey KB, Keefe D, Kourtz L, & Storey JM (1991) Glucose-6-phosphate dehydrogenase in cold hardy insects: kinetic properties, freezing stabilization, and control of hexose monophosphate shunt activity. Insect Biochem. 21(2):157–164.

36. Lippert K & Galinski EA (1992) Enzyme stabilization be ectoine-type compatible solutes: protection against heating, freezing and drying. Appl. Microbiol. Biotechnol. 37(1):61–65.

37. Fabrie CH, de Kruijff B, & de Gier J (1990) Protection by sugars against phase transition-induced leak in hydrated dimyristoylphosphatidylcholine liposomes. Biochimica et Biophysica Acta (BBA)-Biomembranes 1024(2):380–384.

38. Rudolph AS, Crowe JH, & Crowe LM (1986) Effects of three stabilizing agents—proline, betaine, and trehalose—on membrane phospholipids. Arch. Biochem. Biophys. 245(1):134–143.

39. Koštál V, Mollaei M, & Schöttner K (2016) Diapause induction as an interplay between seasonal token stimuli, and modifying and directly limiting factors: hibernation in Chymomyza costata. Physiol. Entomol. 41(4):344–357.

40. Moon I, Fujikawa S, & Shimada K (1996) Cryopreservation of Chymomyza larvae (Diptera: Drosophilidae) at-196°C with extracellular freezing. Cryo-Letters 17:105–110.

41. Shimada K & Riihimaa A (1988) Cold acclimation, inoculative freezing and slow cooling: essential factors contributing to the freezing-tolerance in diapausing larvae of Chymomyza costata (Diptera: Drosophilidae). Cryo Letters 9:5–10.

42. Koštál V, Zahradníčková H, & Šimek P (2011) Hyperprolinemic larvae of the drosophilid fly, Chymomyza costata, survive cryopreservation in liquid nitrogen. Proc. Natl. Acad. Sci. U.S.A. 108(32):13041–13046.

43. Des Marteaux LE, Hůla P, & Koštál V (2019) Transcriptional analysis of insect extreme freeze tolerance. Proc. R. Soc. B 286(1913):20192019.

44. Štětina T, Des Marteaux L, & Koštál V (2020) Insect mitochondria as targets of freezing-induced injury. Proc. R. Soc. B 287(1931):20201273.

45. Tamiya T, et al. (1985) Freeze denaturation of enzymes and its prevention with additives. Cryobiology 22(5):446–456.

46. Hart R, Ramazzotto LJ, & Engstrom R (1972) Cryoprotection of some rat heart enzymes. Cryobiology 9(5):461–464.

47. Yamanaka H & Mackie IM (1971) Changes in activity of a sarcoplasmic adenosinetriphosphatase during iced-storage and frozen-storage of cod. Bull. Jap. Soc. Sci. Fish. 37(11):1105.

48. Connell J (1966) Changes in aldolase activity in cod and haddock during frozen storage. J. Food Sci. 31(3):313–316.

49. Chebotareva N, Kurganov B, & Livanova N (2004) Biochemical effects of molecular crowding. Biochemistry (Moscow) 69(11):1239–1251.

50. Vöpel T & Makhatadze GI (2012) Enzyme activity in the crowded milieu. PLoS One 7(6):e39418.

51. Fiorini E, Börner R, & Sigel RK (2015) Mimicking the in vivo environment–The effect of crowding on RNA and biomacromolecular folding and activity. CHIMIA Int. J. Chem.

52. Drobnis EZ, et al. (1993) Cold shock damage is due to lipid phase transitions in cell membranes: a demonstration using sperm as a model. J. Exp. Zool. 265(4):432–437.

53. Powell D, Sato JD, Brock HW, & Roberts DB (1984) Regulation of synthesis of the larval serum proteins of Drosophila melanogaster. Dev. Biol. 102(1):206–215.

54. Telfer WH & Kunkel JG (1991) The function and evolution of insect storage hexamers. Annu. Rev. Entomol. 36(1):205–228.

55. Moos M, et al. (2022) Cryoprotective metabolites are sourced from both external diet and internal macromolecular reserves during metabolic reprogramming for freeze tolerance in drosophilid fly, Chymomyza costata. Metabolites 12(2):163.

56. Hidalgo M, et al. (2018) Concentrations of non-permeable cryoprotectants and equilibration temperatures are key factors for stallion sperm vitrification success. Anim. Reprod. Sci. 196:91–98.

57. Oldenhof H, et al. (2013) Osmotic stress and membrane phase changes during freezing of stallion sperm: mode of action of cryoprotective agents. Biol. Reprod. 88(3):68, 61-11.

58. Hornberger K, Li R, Duarte ARC, & Hubel A (2021) Natural deep eutectic systems for nature-inspired cryopreservation of cells. AICHE J. 67(2):e17085.

59. Cabrita E, Anel L, & Herraez M (2001) Effect of external cryoprotectants as membrane stabilizers on cryopreserved rainbow trout sperm. Theriogenology 56(4):623–635.

60. Lee RE, McGrath JJ, Morason RT, & Taddeo RM (1993) Survival of intracellular freezing, lipid coalescence and osmotic fragility in fat body cells of the freeze-tolerant gall fly Eurosta solidaginis. J. Insect Physiol. 39(5):445–450.

