## Supplementary Information for "Insect cell plasma membranes do, while soluble enzymes do not, need stabilization by accumulated cryoprotectant molecules during freezing stress"

#### SI Supplementary Methods

**Rearing and acclimation of experimental insects**

A colony of *C. costata*, Sapporo strain (1), was reared on artificial diet in MIR 154 incubators (Sanyo Electric, Osaka, Japan) as described previously (2). Two phenotypic variants were generated according to our earlier acclimation protocols (3): (i) active, warm-acclimated, freeze-sensitive larvae (abbreviated as LD in earlier papers); (ii) diapause, cold-acclimated, freeze-tolerant larvae (abbreviated as SDA in earlier papers). Briefly, the active larvae were reared from eggs to 3rd larval instars (age of 3 weeks) at 18°C under long-day photoperiod (16h light : 8h dark), which allows direct non-diapause development to pupa and adult. The diapause larvae were reared until age of 6 weeks at 18°C under short-day photoperiod (12h light : 12h dark), which induces larval diapause; next, diapause larvae were transferred to constant darkness and progressively cold acclimated over five weeks (1 week at 11°C, followed by 4 weeks at 4°C).

The active larvae of *C. costata* have limited survival after freezing stress (35% survive slow inoculative freezing to -5°C, 10% to -10°C, and none survive freezing to -20°C or below). In contrast, practically all diapause larvae survive freezing to -30°C or even -75°C, and 42.5% survive and metamorphose into fit adults after 18 months of cryopreservation in $LN_2$ (4). Since we focused on the nature of damage caused by freezing stress, the freeze-sensitive, active larvae were used for most experiments. The diapause larvae were used for the experiments on integrity of plasma membrane upon freezing stress.

For supplementary experiments, we used larvae of vinegar fly, *Drososphila melanogaster* (Oregon R strain) and adult locusts, *Locusta* migratoria coming from insect cultures routinely maintained in Biology Centre CAS, České Budějovice, Czech Republic. The vinegar flies were reared at constant 18°C, while locusts were maintained at constant 30°C, both species were kept at a photoperiodic regime 12h ligth : 12h darkness. Both species represented warm-adapted and warm-acclimated animals with relatively low cold tolerance, killed by chilling to temperatures around 0°C for minutes or hours (i.e. without freezing) (5, 6).

#### General outline of experiments

The general outline of all experiments is presented in Fig. S1. The experiments were performed in two different ways: *in vivo* and *in vitro*.

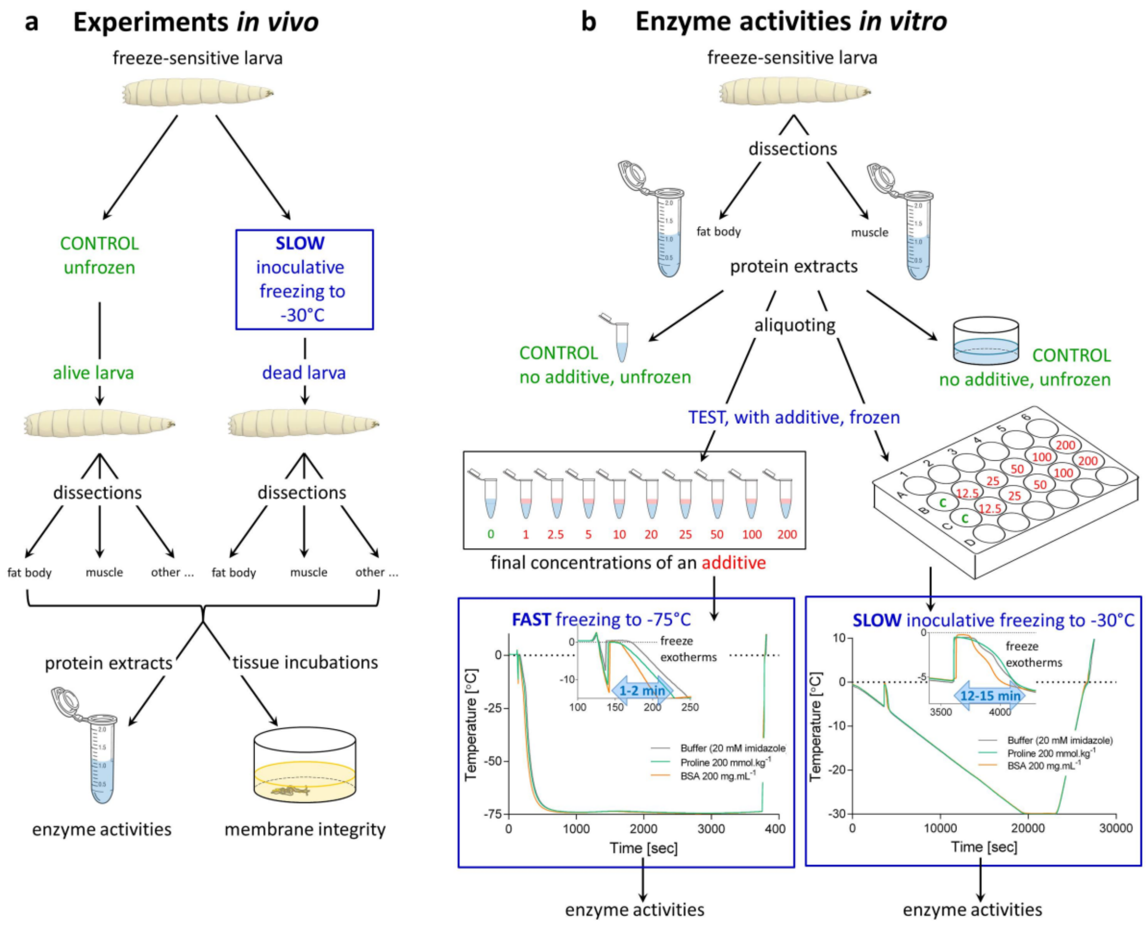


**Fig. S1: The schematic outline of experiments.**

**(a) Experiments *in vivo*.** Whole larvae of the freeze-sensitive phenotype of *Chymomyza costata* were exposed to lethal slow inoculative freezing to -30°C using the protocol described earlier (4). Unfrozen larvae served as controls. Upon melting, (i) different tissues of larvae were rapidly dissected, crude total proteins were extracted, which served as source for analysis of enzyme activities, or (ii) 'intestine blobs' were rapidly dissected, incubated in Trypan Blue stain, and membrane integrity was assessed. For more explanations, see text (section *In vivo* experiments).

**(b) Enzyme activities *in vitro*.** The fat body or muscle tissues were dissected from freeze-sensitive larvae and total proteins were extracted in 20 mM imidazole buffer, pH 7.2. The extract was aliquoted and the aliquots were exposed to freezing in the absence or presence of different additives. One aliquot served as control – no additive and not frozen. The enzyme activities were measured upon melting. We used two different freezing protocols: (i) FAST freezing to -75°C; and (ii) SLOW inoculative freezing to -30°C (same protocol as used for whole larvae in the *in vivo* experiments). The examples of temperature courses in the FAST and SLOW protocols are shown in blue frames for plain buffer (20 mM imidazole), buffer augmented with proline (200 $mmol.kg^{-1}$), and buffer augmented with BSA (200$mg.mL^{-1}$). The records were obtained using K-type thermocouples attached to PicoLog TC-08 datalogger (Pico technology, St. Neots, UK). Note that additives had relatively small influence on the course of temperature and that the durations of freeze-exotherms was longer in the SLOW than in the FAST protocol (blue arrows in the insets). For more explanations, see text (section *In vitro* enzyme activity assays).

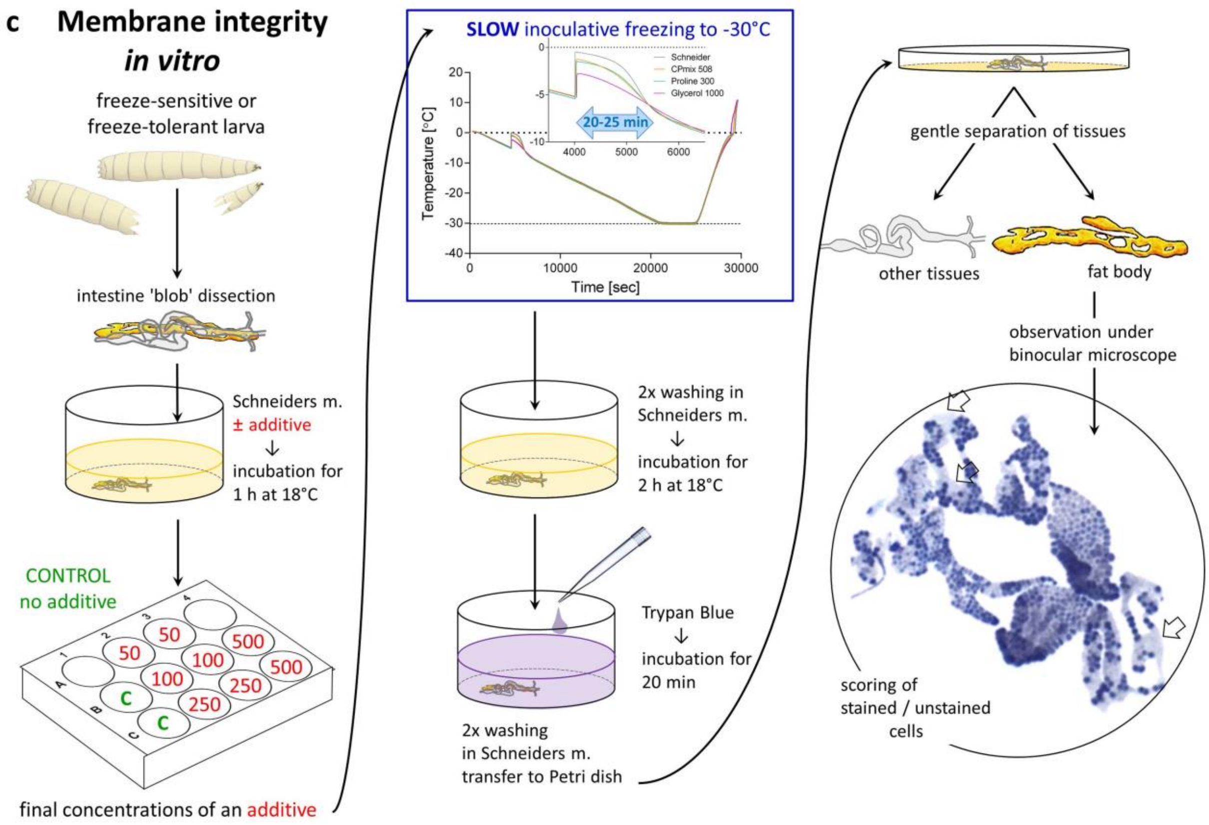


**Fig. S1 - continues: The schematic outline of experiments.**

**(c) Membrane integrity *in vitro*.** The 'intestine blobs' were dissected from larvae of the freeze-sensitive or freeze-tolerant phenotypes of *Chymomyza costata* and pre-incubated for 1h in Schneider's medium in the absence or presence of different additives. Next, the intestines were exposed to SLOW inoculative freezing to -30°C (same protocol as used for whole larvae in the *in vivo* experiments). Upon melting, the additives were washed out and the intestines were incubated in Trypan Blue solution. Next, the fat body tissues were gently separated from 'intestine blobs', observed under binocular microscope, and the proportion of blue-stained cells was estimated. The examples of temperature courses during SLOW freezing are shown in blue frame for plain Schneider's medium, and for Schneider's medium augmented by: CPmix 508 (a mixture of *C. costata* native cryoprotectants at concentrations corresponding to hemolymph of freeze-tolerant larva, sum concentration of 508 $mmol.kg^{-1}$), proline (300 $mmol.kg^{-1}$), and glycerol (1 000 $mmol.kg^{-1}$). The records were obtained using K-type thermocouples attached to PicoLog TC-08 datalogger (Pico technology, St. Neots, UK). Note, that additives had relatively small influence on the course of temperature and that the durations of freeze-exotherms was approximately 20-25 min. For more explanations, see text (section *In vitro* membrane integrity asssays).

**Table S1: The list of additives**

| Additive | Sigma-Aldrich product | | m.w. | Medium | Concentration in plain medium [mg/mL] * | Amount in mg added to 1 mL of plain medium to produce given concentration ** | |
|---|---|---|---|---|---|---|---|
| | Name | Number | | | | | |
| **Proline** | L-Proline | P0380 | 115.13 | 20 mM imidazole | 0 | 11.5 | 100 $mmol.kg^{-1}$ |
| | | | | Schneider's | 1.70 | 9.8 | 100 $mmol.kg^{-1}$ |
| **Trehalose** | D(+) Trehalose dihydrate | T9449 | 378.33 | 20 mM imidazole | 0 | 37.8 | 100 $mmol.kg^{-1}$ |
| | | | | Schneider's | 2.0 | 35.8 | 100 $mmol.kg^{-1}$ |
| **Glutamine** | L-Glutamine | G3126 | 146.14 | 20 mM imidazole | 0 | 14.6 | 100 $mmol.kg^{-1}$ |
| | | | | Schneider's | 1.8 | 12.8 | 100 $mmol.kg^{-1}$ |
| **Asparagine** | L-Asparagine | A0884 | 132.12 | 20 mM imidazole s | 0 | 13.2 | 100 $mmol.kg^{-1}$ |
| | | | | Schneider's | 0 | 13.2 | 100 $mmol.kg^{-1}$ |
| **Betaine** | Betaine | 61962 | 117.15 | 20 mM imidazole | 0 | 11.7 | 100 $mmol.kg^{-1}$ |
| | | | | Schneider's | 0 | 11.7 | 100 $mmol.kg^{-1}$ |
| **Glycerol** | Glycerol | G5516 | 92.09 | 20 mM imidazole | 0 | 9.2 | 100 $mmol.kg^{-1}$ |
| **Saccharose** | Sucrose | S1888 | 342.30 | Schneider's | 0 | 34.2 | 100 $mmol.kg^{-1}$ |
| **Histodenz** | Histodenz | 1D2158 | 821.14 | 20 mM imidazole | 0 | 82.1 | 100 $mmol.kg^{-1}$ |
| | | | | Schneider's | 0 | 82.1 | 100 $mmol.kg^{-1}$ |
| **Urea** | Urea | U5378 | 60.06 | 20 mM imidazole | 0 | 6.0 | 100 $mmol.kg^{-1}$ |
| **$K^+$** | Potassium chloride | P5405 | 74.56 | 20 mM imidazole | 0 | 7.5 | 100 $mmol.kg^{-1}$ |
| **$Na^+$** | Sodium chloride | S5886 | 58.44 | Schneider's £ | 1.16 | 1.14 ($Na^+$)<br>2.87 (NaCl) | 100 $mmol.kg^{-1}$ |
| **BSA** | Bovine Serum Albumin | A7030 | 66 430 | 20 mM imidazole | 0 | 100.0 | 100 $mg.mL^{-1}$ |
| | | | | Schneider's $ | 0 | 100.0 | 100 $mg.mL^{-1}$ |
| **Ficoll** | Ficoll PM 70 | F2878 | ca. 70 000 | 20 mM imidazole | 0 | 100.0 | 100 $mg.mL^{-1}$ |
| | | | | Schneider's | 0 | 100.0 | 100 $mg.mL^{-1}$ |

* no 'additives' were present in 20 mM imidazole buffer, while some 'additives' are present in the Schneider's *Drosophila* medium and those were taken into account when preparing the augmented media. The concentration data are according to Biosera formulation of 21/07/2017.

** this is an example showing amounts of additives used to prepare augmented imidazole buffer (for Enzyme experiments) or Schneider's medium (for Membrane experiments) of a given concentration. For simplicity, we considered the mass of 1 mL of 20 mM imidazole or 1 mL of Schneider's medium to be 1 g.

£ the sodium ion ($Na^+$) is present in the Schneider's medium in three forms (NaCl, $NaHCO_3$, and $Na_2HPO_4$), which in sum contribute 1.16 mg of sodium ion per mL. We used NaCl to augment the Schneiders's medium by sodium ions.

$ the Schneider's solution contains yeast extract ( 2g/L) which brings some additional protein (not taken into account).

***In vivo* experiments**

In the *in vivo* experiments (Fig. S1a), whole larvae were exposed to slow inoculative freezing to -30°C using the protocol described earlier (4). The protocol consisted of five steps set in a Ministat 240 programmable cryostat (Huber, Offenburg, Germany): (i) 20 min of larval manipulation at 0°C (washing larvae out of the diet, dividing into groups of 20 specimens, and placing them into 3 mL plastic tubes between two layers of moist cellulose); (ii) slow pre-freezing to -30°C (with a small ice crystal added on top of the moist cellulose) for 300 min (cooling rate, 0.1°C $min^{-1}$); (iii) keeping larvae at -30°C for 60 min; (iv) heating from -30°C to +5°C over 60 min (heating rate, 0.6°C $min^{-1}$); and (v) melting at constant +5°C for 10 min. Adding a small ice crystal at the beginning of step (ii) induces inoculative internal freezing in larvae, which starts between -1 and -2°C, the formation of ice fraction (76% of total body water is freezable in active larvae) is completed at -5.9°C (i.e. takes approximately 59 min) (4). Unfrozen larvae served as controls. Right upon melting (end of step v): either different tissues of larvae were rapidly dissected, crude total proteins were extracted, which served as source for analysis of enzyme activities, or 'intestine blobs' were rapidly dissected, incubated in Trypan Blue stain, and fat body plasma membrane integrity was assessed. The enzyme activity and membrane integrity assays are described later in the text.

***In vitro* enzyme activity assays**

To conduct *in vitro* assays of enzyme activities (Fig. S1b), the fat body or muscle tissues were rapidly dissected (ca. 30 sec per one larva) from 30 freeze-sensitive larvae and pooled in 1200 µL of ice cold 20 mM imidazole buffer, pH 7.2. Extracting 30 tissues in 1200 uL of buffer, we estimate that the tissue solutions were diluted approximately 200-fold, which means that they could contribute additional 2.3 mOsm concentration of tissue solutes to the sample (the larval hemolymph has an osmotic concentration of 465 mOsm according to (7)).

The pools of dissected tissues were homogenized for 6 x 3 sec using X120 homogenizer with metal blades (CAT, Germany), centrifuged at 21,000 g for 10 min at 4°C, and the supernatant was used as the source of enzymes. The supernatant was aliquoted; one aliquot served as control ─ no additive and not frozen ─ and was used for enzyme activity assay after incubation on ice for 1h (control = 100% activity). The other aliquots were exposed to freezing in the absence or presence of different additives (see Table S1 for complete list of additives) and the enzyme activities were measured upon melting. We used two different freezing protocols: (i) FAST, which simulates routine laboratory practice when a protein extract in a plastic micro-vial is simply moved to a deep freezer set to -75°C; and (ii) SLOW, which simulates ecologically relevant, slow inoculative freezing as used for whole larvae (*in vivo* experiments).

For the FAST freezing, we used 50 uL aliquots of protein extract mixed with 20 uL volume of additive solution 20 mM imidazole (when no additive was used, 20 uL of plain 20 mM imidazole was added). Mixing the extract and additive solution in 200 uL PCR tubes, different concentrations of additives were produced ranging between 0 and 200 $mmol.kg^{-1}$ (or 0 and 200 $mg.mL^{-1}$ for macromolecular compounds BSA and Ficoll). The PCR tubes were moved to deep freezer (Platinum 370H, Angelantoni, Italy) for 1h, then melted on ice and immediately used for enzyme activity assays. The freeze exotherms started after spontaneous formation of ice nucleus (at temperatures close to -10°C) and were relatively short (1-2 min) (see inset in FAST freezing protocol in Fig. S1b).

For the SLOW inoculative freezing, we used 100 uL aliquots of protein extract mixed with 100 uL of 20 mM imidazole in the well of 24-well polysterene Costar plate with untreated surface (Corning, Kennebunk, ME, USA). In this total volume of 200 uL, given amount of additives were dissolved to produce concentations ranging between 0 and 500 mmol.kg$^{-1}$ (or 0 and 200 mg.mL$^{-1}$ for macromolecular compounds BSA and Ficoll). The plate was moved to Ministat 240 (Huber) and the same protocol as used for whole larvae was started. Upon melting, the enzyme activities were assessed. The ice formation was nucleated by adding external ice crystal (frozen 25 uL droplet of 20 mM imidazole) at exactly -5°C and the freeze exotherms lasted for 12-15 min (see inset in SLOW freezing protocol in Fig. S1b). Exact methods to measure the enzyme activities are described below (section: Enzyme activity assays).

***In vitro* membrane integrity assays**

To conduct *in vitro* assays on plasma membrane integrity (Fig. S1c), we dissected the 'intestine blobs' from larvae under Schneider's *Drosophila* medium (Biosera, Nuaillé, France), which is widely used to support viability of incubated insect tissues. The intestines were obtained by cutting off the front tip of larval body and gently squeezing the intestines out of the integument. No further attempt was made to separate individual tissues from the blob as this would result in damaging many cells. The whole intestine blobs were moved to 650 μL of Schneider's medium in the wells of 12-well tissue-culture plate (TRP, Switzerland). Typically, ten blobs were placed in one well though this number varied between six and twelve according to availability of larvae. The Schneider's medium has a complex composition including different metabolites, salts and buffers (total osmolarity of 330 mOsm). We changed this composition by adding additives (Table S1) at different concentrations ranging between 0 and 1000 mmol.kg$^{-1}$ (or 0 and 200 mg.mL$^{-1}$ for macromolecular compounds BSA and Ficoll). The final concentrations were prepared by dissolving the given amount of additive in a known small volume of Schneider's medium. That is why we express the concentrations in molal units. For simplicity, we considered the mass of 1L of the Schneider's solution to be 1kg. All concentrations we refer to are the final concentrations in the augmented medium taking into account the concentrations of respective compound in the original Schneider's medium. No additives were used in controls.

The intestines were pre-incubated in their respective medium at 18°C for 1h prior to the freezing stress. The 12-well-design of the plate allowed variously combining controls, treatments and their replicates; two wells were always reserved for temperature record (no tissues). One well was taken as one replication, i.e. the proportion of blue-stained cells was estimated for all fat body tissues in the well (six to twelve) and the mean blue staining of the group was taken for calculation. Each treatment (i.e. a combination of the additive and concentration) was replicated at least twice (two wells) but some treatments were replicated 3-7 times and the control was replicated 11 times (the exact numbers of replicates are shown in respective Figures). After the pre-incubation, whole plate was housed in the Ministat 240 (Huber) and the same protocol as used for whole larvae was started. The ice formation was nucleated by adding external ice crystal (frozen 50 uL droplet of Schneider's medium) at exactly -5°C and the freeze exotherms lasted for

20-25 min (see inset in SLOW freezing protocol in Fig. S1c). Further processing of intestine blobs after melting is described below (section: Trypan Blue assay of plasma membrane integrity).

**Enzyme activity assays**

The activities of seven different enzymes were measured according to protocols specified below. The activities were measured colorimetrically (most enzymes) or fluorimetrically (*matrix metalloproteinases*) using either: (i) Quartz glass cuvettes housed in the Cary50 UV-Vis spectrophotometer (Varian, USA) attached to circulating water bath BC4 (Julabo, Germany) for controlling the constant 25°C in cuvettes, or (ii) 96-well Optical Btm PolymerBase plates (ThermoScientific, USA) housed in the Infinite 200Pro plate reader (Tecan, Austria) equilibrated to constant 25°C (most enzymes) or 30°C (*phenoloxidase*). All activities were measured in a kinetic mode during the initial (linear) part of enzyme kinetics. We continuously monitored the change in the absorbance/fluorescence intensity over time, which is linked to consumption/production of a specific substrate/product. All chemicals used for enzyme activity assays were either included in the respective enzyme assay kit, or were purchased from Sigma-Aldrich.

For the *in vivo* experiments, three replicates of unfrozen control larvae vs. three replicates of slowly frozen larvae were compared and each replicate was a pool of group of larvae or tissues (as specified below). The composition of extraction buffers and reaction mixtures are specified below for each enzyme. The enzyme activities were expressed in micromoles of enzymatically converted substrates per min per mg of total protein (except for *phenoloxidase* and *matrix metalloproteinases*, see below). The total proteins were measured by bicinchoninic acid assay (BCA) according to (8).

For the *in vitro* experiments, single large-volume extract of 30 larval tissues was prepared by homogenization in 20 mM imidazole, pH 7.2. This extract was divided into equal aliquots, the aliquots exposed to freezing (except control), and the enzyme activities were measured: (i) the activity in unfrozen control was taken as 100%; (ii) the other aliquots were frozen in presence of various additives at concentrations ranging between 0 and 500 $mmol.kg^{-1}$. The enzyme activities were expressed as a fraction (in percentage) of enzyme activity that recovers after freezing stress: we searched for the additive concentration ($P_{50}$) that ensures 50% recovery of initial (unfrozen control) enzyme activity.

*Glucose 6-phosphate dehydrogenase* (G6PDH) activity was measured in total protein extracts from larval fat body or larval muscles using Cary50 UV-Vis spectrophotometer. Pools of 30 tissues larvae per replicate were homogenized in 400 µL of ice cold 100 mM Tris-HCl buffer, pH 8.0, containing 15 mM mercaptoethanol, and 1 mM EDTA (600 µL of 20 mM imidazole buffer was used for *in vitro* assays). The composition of reaction mixture for G6PDH assay was same as described earlier (9): 20 mM imidazole-HCl buffer, pH 7.2; 5 mM $MgSO_4$; 0.6 mM $NADP^+$; 1 mM glucose 6-phosphate; and larval protein extract (50 uL in a total of 500 uL of reaction mixture). The reaction was initiated by the addition of of glucose 6-phosphate solution into reaction mixture and its conversion to 6-phosphate gluconolactone by G6PDH activity was followed by measuring the increase of 340 nm absorbance ($NADP^+$ → NADPH).

*Citrate synthase* (CS) activity was measured in total protein extracts from larval fat body or larval muscles using Cary50 UV-Vis spectrophotometer. Pools of 30 tissues per replicate were homogenized in 400 µL of ice cold in 50 mM Tris buffer, pH 8.0, containing 0.2 M sucrose, 1 mM EDTA, and 1% Triton X (600 µL of 20 mM imidazole buffer was used for *in vitro* assays). In addition to standardly homogenizing the tissues using CAT homogenizer, we sonicated the samples 5 times for 1 sec (4710 Series Ultrasonic Homogenizer, Cole Parmer, Chicago. IL, USA) in order to solubilize the enzyme from mitochondrial matrix. The composition of reaction mixture for CS assay was same as described earlier (10): 50 mM imidazole-HCl buffer, pH 8.0; 0.05 mM DTNB reagent (5,5'-dithiobis-2-nitrobenzoic acid); 0.15 mM acetyl CoA; 0.25 mM oxaloacetate; and larval protein extract (10 uL in a total of 500 uL of reaction mixture). The reaction was initiated by the addition of 25 µL of oxaloacetate solution into 475 µL of reaction mixture and the mercaptid ion formation from CoA-SH (that was released upon synthesis of citrate) was followed by measuring the increase of 412 nm absorbance (DTNB → TNB).

*Lactate dehydrogenase* (LDH) activity was measured in total protein extracts from larval muscles of *C. costata* using Cary50 UV-Vis spectrophotometer. Pools of 30 muscle tissues per replicate were homogenized in 400 µL of ice cold 100 mM Tris-HCl buffer, pH 8.0, containing 15 mM mercaptoethanol, and 1 mM EDTA (600 µL of 20 mM imidazole buffer was used for *in vitro* assays). In addition to *C. costata*, we measured the activity of LDH (using the same method) also in the muscles of the 3rd instar larvae of *Drosophila melanogaster* (pool of 30 tissues per replicate), and in the tibial levator muscle dissected from femur of the hind leg of the adult locust, *Locusta migratoria* (single muscle per replicate). Both insects were taken from our routine cultures at the Biology Centre, České Budějovice, and were acclimated to warm conditions (*D. melanogaster* is reared at constant 18°C, while *L. migratoria* at constant 30°C). The composition of reaction mixture for CS assay was same as described earlier (11): 80 mM Tris-HCl, pH 7.5; 100 mM KCl ; 0.2 mM NADH; 2 mM pyruvate. The reaction was initiated by the addition of 10 µL of protein extract into 490 µL of reaction mixture and the LDH activity was followed by measuring the decrease of 340 nm absorbance (NADH → $NAD^+$).

*Phenoloxidase* (PO) activity was measured in hemolymph using Infinite 200Pro plate reader. The hemolymph was obtained by gently tearing of larvae on a piece of Parafilm, creating a large droplet of pooled hemolymph (pool of 30 larvae was bled per replicate). This droplet was then extracted using a calibrated glass capillary (Drummond Sci., Broomall, PA, USA). The PO activity was assayed through its catalytic conversion of L-dopa (3,4-dihydroxy-L-phenylalanine, colourless) to dopachrome (red-brown colour) according to (12): the following components were mixed in the well of 96-well plate: 5 µL of hemolymph; 35 µL of water; 5 µL of PBS, and 20 µL of 5 mM L-DOPA solution. The plate was moved to plate reader equilibrated to 30°C, briefly shaked (10 sec), and darkening of the sample was continuously followed as the increase of 490 nm absorbance (L-DOPA → dopachrome).

General activity of *matrix metalloproteinases* (MMPs, collagenases) was measured in total protein extracts from whole larval homogenates using Infinite 200Pro plate reader according to instructions for use of the MMP Activity Assay Kit (ab112147, Abcam, Cambridge, UK). Pools of ten larvae per replicate were homogenized in 500 µL of ice cold 100 mM Tris-HCl buffer, pH 7.2. The 25 µL aliquot of the supernatant from centrifugation was used as a source of enzyme, combined with 25 µL of 2 mM APMA (4-aminophenylmercuric acetate) in the well of 96-well plate, and incubated for 15 min at room temperature. Next, 50 µL of the artificial Red MMP Substrate (FRET peptide) diluted in Assay Buffer (1:100) was added to the well and whole plate was moved to the plate reader. Upon cleavage of the FRET peptide by MMPs, the intensity of fluorescence at 540/590 nm (Ex/Em) increases, which was monitored over time (RFU, relative fluorescence units).

General activity of *amylases and maltases* was measured in total protein extracts from larval midguts using Infinite 200Pro plate reader according to instructions for use of the Amylase Assay Kit (ab102523, Abcam). Pools of ten larval midguts per replicate were homogenized in 400 µL of the Assay Buffer. The 50 µL aliquot of the supernatant from centrifugation was used as a source of enzyme and combined with 100 µL of artificial substrate ethylidene-*p*NP-G7 diluted in Assay Buffer (1:1) in the well of 96-well plate. The plate was moved to the plate reader. The α-amylase cleaves the artificial substrate to produce smaller fragments that are eventually modified by maltases, causing the release of a chromophore (nitrophenol) that was measured as the increase of 405 nm absorbance. The enzyme activity was converted to $\mu mol.min^{-1}.mg^{-1}$ total protein using calibration curve produced by measuring absorbances at 405 nm for different concentrations of nitrophenol.

*Glycogen phosphorylase* activity was measured in total protein extracts from larval muscles using Cary50 UV-Vis spectrophotometer. Pools of 20 larval muscles per replicate were homogenized in 400 µL of 100 mM Tris-HCl buffer, pH 8.0 containing 15 mM mercaptoethanol and 1 mM EDTA. The composition of reaction mixture was same as described earlier (13): 50 mM potassium phosphate ($KH_2PO_4$) buffer, pH 6.8; 5 mg $mL^{-1}$ glycogen, 5 µM glucose-1,6-diphosphate, 0.6 mM $NADP^+$, 2 mM 5'AMP, 15 mM $MgCl_2$, 2 U $mL^{-1}$ phosphoglucomutase (PGM), and 2 U $mL^{-1}$ glucose 6-P dehydrogenase (G6PDH). First, the activity of the *active* ('a') form of the enzyme was measured in the absence of 5'AMP. The reaction was initiated by the addition of 70 µL of protein extract into 420 µL of reaction mixture. The glucose units released from glycogen by glycogen phosphorylase activity are converted by coupling enzymes (PGM and G6PDH) to gluconolactone, while $NADP^+$ is oxidized. This conversion was followed by measuring the increase of 340 nm absorbance ($NADP^+ \rightarrow NADPH$). Next, 10 uL of 5'AMP solution was added, which activates the *inactive* ('b') form of the enzyme (14), and the combined activity of ('a+b') forms was measured.

**Trypan Blue assay of plasma membrane integrity**

The integrity of fat body cell plasma membrane was assessed using the Trypan Blue assay (15). Trypan Blue is negatively charged and large molecule that is cell membrane impermeable and therefore only enters cells with damaged or compromised membranes. Upon entry into the cell, Trypan Blue binds to intracellular proteins and DNA thereby rendering the cells a bluish color.

Upon melting of the medium in wells of 12-well plate (see Fig. S1c), the intestine blobs were immediately washed twice using Schneider's medium in order to remove the additives and the intestines were then incubated in Schneider's medium for 2h at 18°C. Next, the Trypan Blue solution was added (400 µL of 1% solution per well) and intestines were incubated in it for 20 min. Next, the intestines were washed twice using Schneider's medium (in order to remove excess of Trypan Blue) and moved to plastic Petri dish (3 cm diam.) with a thin layer of Schneider's medium covering the bottom. The tissues were gently pushed apart to make fat body lobes nicely visible. The cells of fat body tissues were inspected under binocular microscope at 25x magnification and the percentage of blue-stained cells was estimated. One example of fat body tissue (frozen without additives) is shown in Fig. S1c (estimation: 95% cells blue-stained; the arrows indicate areas with unstained cells).

In a preliminary experiment, we verified that various additives themselves (without freezing stress) have practically no effect on membrane integrity (weak blue-staining) when tissues are incubated for long time (18h) at 18°C in the augmented Schneider's medium, while incubation of tissues for short time (2h) at relatively low concentrations of digitonin (0.01 – 1.0 mmol.kg$^{-1}$) caused severe blue-staining in fat body cells (Fig. S8a). Digitonin is a mild detergent that is routinely used to permeabilize plasma membranes of different cells (16).

**Differential scanning calorimetry (DSC)**

We analyzed thermal transitions in the freezing solutions combining CP (proline or trehalose) and perturbant (urea) that were used for SLOW inoculative freezing in the *in vitro* experiments on enzyme stability. The solution (10 µL) was pipetted into the aluminum DSC pan (volume of 50 µL) and its exact amount was checked by weighing the sample with precision to 0.01 mg using Metler Toledo NewClassic MS balance. Hermetically sealed pans were then loaded into Differential Scanning Calorimeter DSC4000 (Perkin Elmer, Waltham, MA, USA) and frozen and thawed according the following protocol: (i) hold for 1 min at 0°C, (ii) cool to −30°C at 2°C.min$^{-1}$, (iii) hold for 5 min at −30°C, and (iv) warm to 25°C at 10°C min$^{-1}$. An empty pan was used as a reference. The resulting heating curves (see Fig. S7a for examples) were analyzed using Pyris Software (Perkin Elmer). The fraction of freezable, i.e. osmotically active water (OAW) was derived from the area under melting endotherm of water using the standard heat of fusion for the ice/water transition of 334.5 J.g$^{-1}$. The OAW fraction served for estimation of the freeze-concentration of urea in different solutions as follows: if the initial concentrations were, for example, 500 mmol.kg$^{-1}$ of urea and 500 mmol.kg$^{-1}$ of proline, the OAW fraction was 81.81% after freezing to -30°C, meaning that the urea was freeze-concentrated 5.5-fold, and reached the concentration of 2749.5 mmol.kg$^{-1}$ in the remaining unfrozen solution (see Table S2 for all results of DSC analyses and calculations of the freeze-concentrations of urea).

### SI Supplementary Results

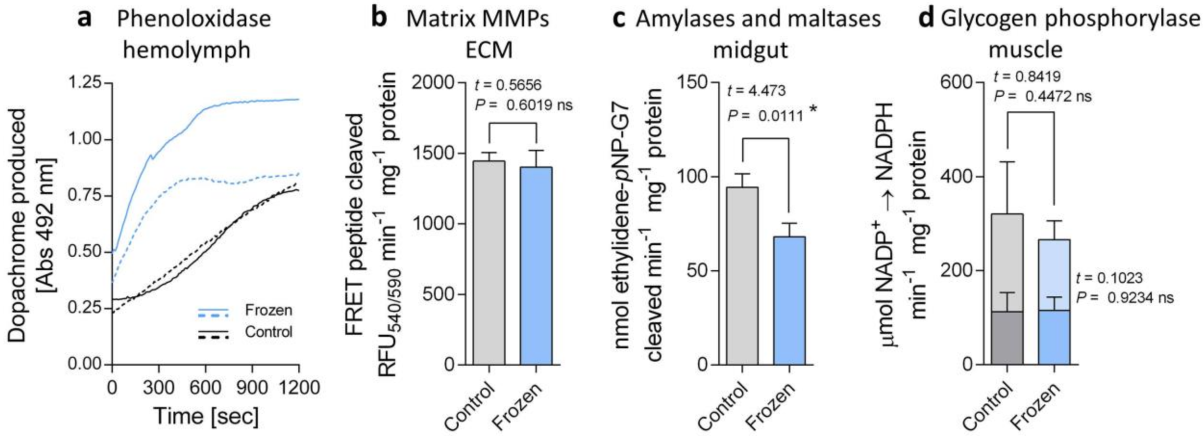


**Fig. S2: The enzyme activities persist after organismally-lethal freezing stress – supplement to Fig. 1.**
The larvae of freeze-sensitive phenotype were killed by freezing to -30°C ('Frozen'). The 'Control' larvae were not frozen. The activities of four different enzymes: (**a**) phenoloxidase; (**b**) matrix metalloproteinases (MMPs); (**c**) amylases and maltases; and (**d**) glycogen phosphorylase were measured in total protein extracts of the respective tissues dissected from Control and Frozen larvae right upon melting. For phenoloxidase (a), we show the time records of the changes in absorbance in two independent replicates (distinguished as solid and broken lines), each replicate represents a pool of hemolymph sampled from 30 larvae. In both replicates, the activity of phenoloxidase in hemolymph of Frozen larvae was relatively high during first 10 min of analysis and then ceased (no further increase of Abs492 was observed), while it was relatively low in Control larvae and a tendency to cessation was seen in one of two replicates (black, solid line). For glycogen phosphorylase (d), we distinguished between active form ('a', lower portions of bars) and inactive form ('b', upper portions of bars) that was activated by 5'AMP. In (b, c, d), each column shows the mean + S.D. of the enzyme activity ($n$ = 3, each replicate represents a pool of 30 larval tissues). The enzyme activities in Control vs Frozen larvae were compared using unpaired, two-tailed $t$ test ($t$ statistics and $P$ values are shown: ns, not significant; * significant difference; GraphPad Prism v. 6.07).

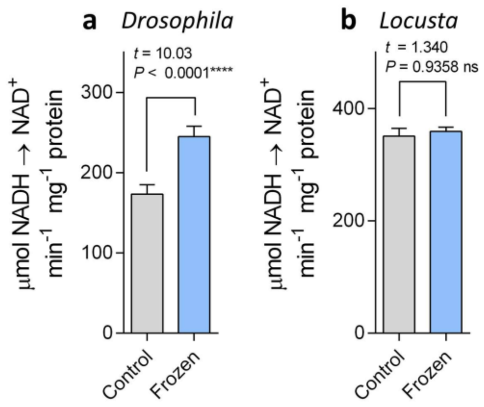


**Fig. S3: The LDH activity after *in vivo* freezing of larvae of *Drosophila melanogaster* and femoral muscle of *Locusta migratoria*.**

The larvae of *D. melanogaster* (**a**) and femoral muscle of *L. migratoria* (**b**) were frozen to -30°C ('Frozen'). The 'Control' specimens were not frozen. The activity of lactate dehydrogenase (LDH) was measured in total protein extracts from muscles obtained right upon melting. Each column shows the mean + S.D. of the enzyme activity ($n$ = 3, each replicate represents a pool of 30 larval tissues (a) or single femoral muscle (b)). The enzyme activities in Control vs Frozen larvae were compared using unpaired, two-tailed $t$ test ($t$ statistics and $P$ values are shown: ns, not significant; **** significant difference; GraphPad Prism v. 6.07).

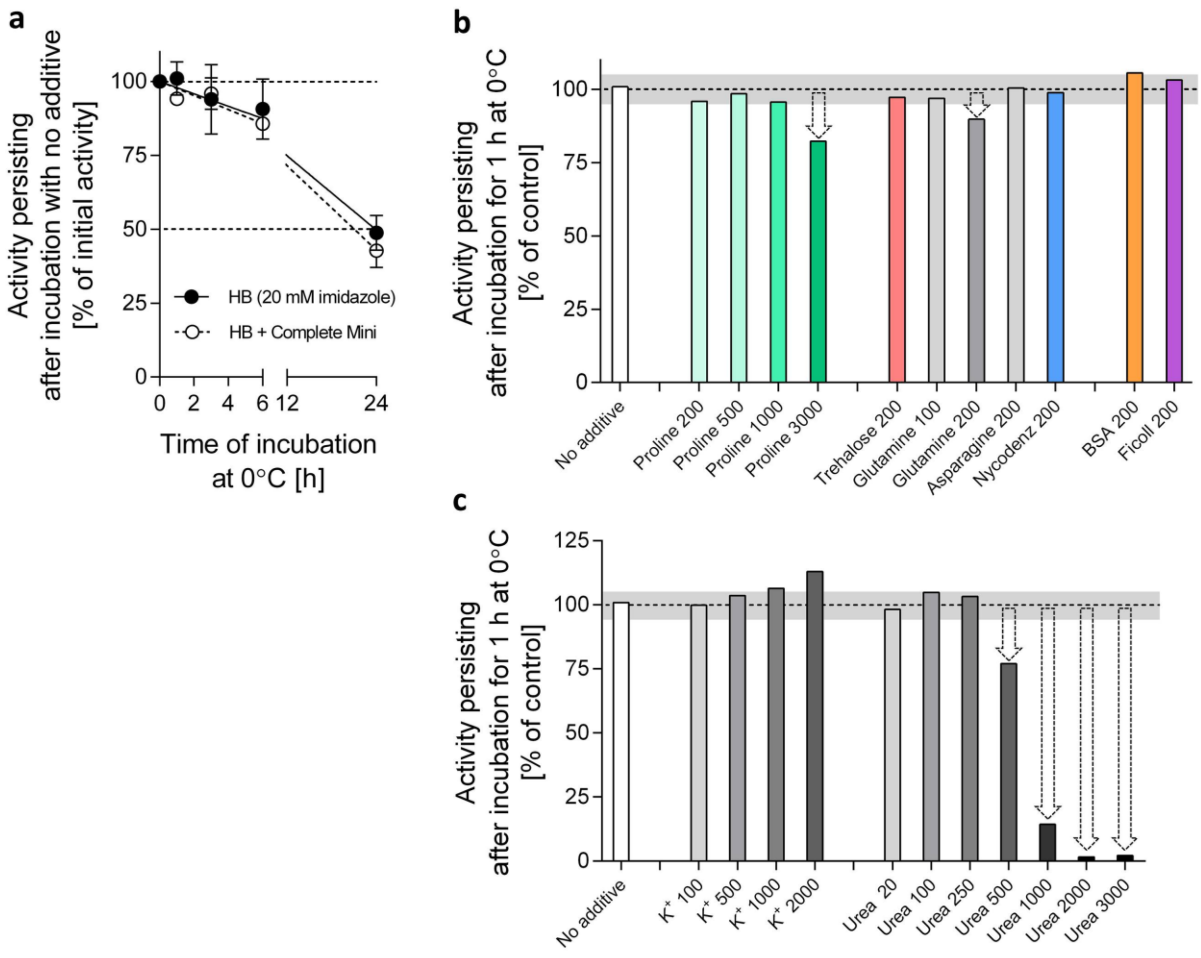


**Fig. S4: The effect of incubation at 0°C with different additives on activity of G6PDH extracted from larval muscles of *Chymomyza costata*.**

The total proteins were extracted in homogenization buffer (HB, 20 mM imidazole) from the fat body tissue of freeze-sensitive phenotype of *Chymomyza costata* larvae and the activity of glucose 6-phosphate dehydrogenase (G6PDH) was measured. (**a**) The extract aliquots were incubated at 0°C for 0–24 h and the gradual loss of activity over time of incubation was registered. Note that the extraction of proteins with or without the proteinase inhibitor cocktail (Complete Mini, EDTA free, Roche, Mannheim, Germany) had no effect on enzyme activity. Each point is a mean ± SD of five replicates. The lines are linear regressions: HB, $y$ = -2.096*x+100; HB+Complete Mini, $y$ = -2.377*$x$+100. (**b, c**) Different additives were added to HB at concentrations indicated and their effect on G6PDH activity after 1h-incubation at 0°C was assessed. High concentrations of proline (3000 mmol.kg$^{-1}$) and glutamine (200 mmol.kg$^{-1}$) suggested moderate decrease of enzyme activity (arrows) below the control level [dashed line is a mean and grey field delimits ± SD of enzyme activity in the control without additives, see panel (a)]. Urea administered at concentrations > 500 mmol.kg$^{-1}$ caused drastic decrease of the enzyme activity. Each column is one replicate except for control [see panel (a)].

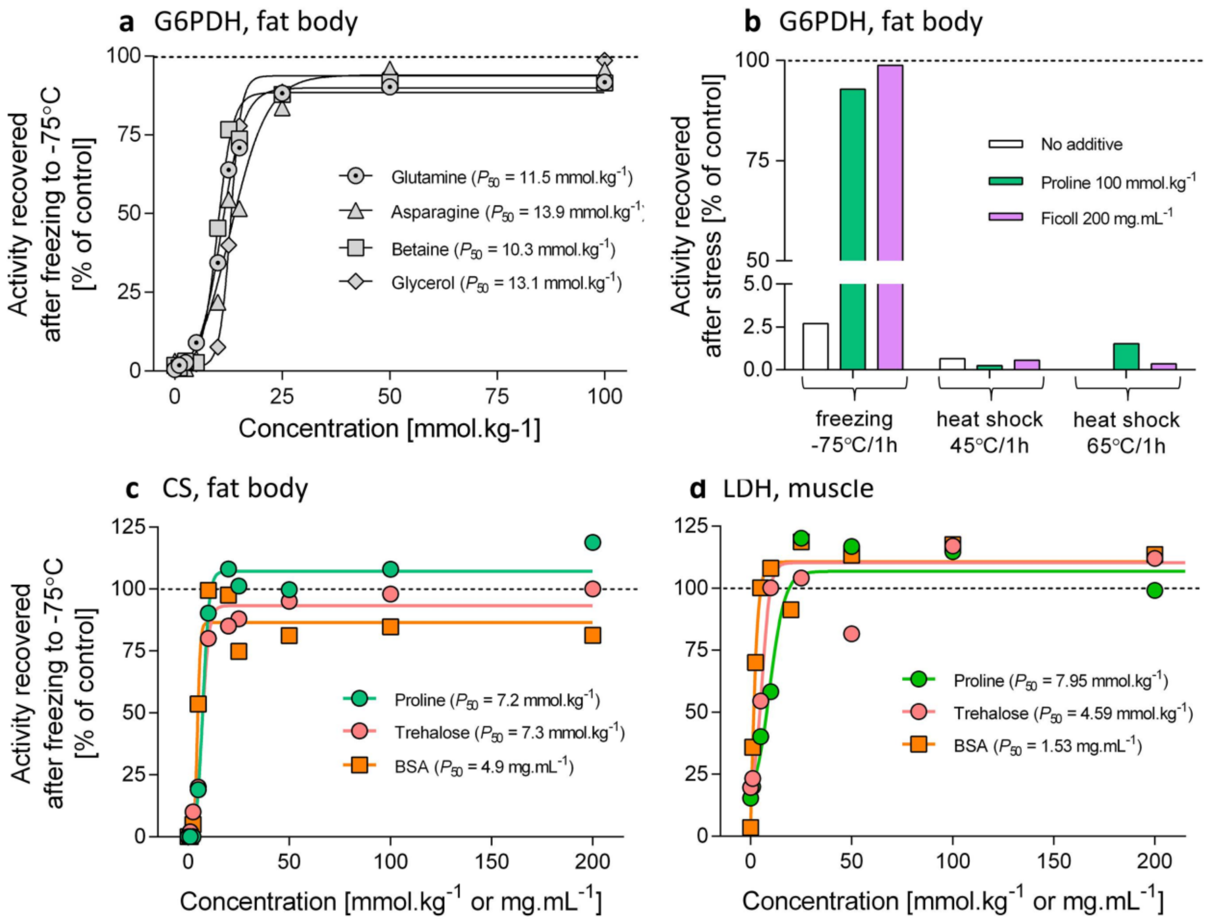


**Fig. S5: The enzyme activity is protected by low concentrations of various additives to the *in vitro* freezing solution – supplement to Fig 2.**

The total proteins were extracted from the fat body or muscle tissues of freeze-sensitive phenotype of *Chymomyza costata* larvae and the activity of glucose 6-phosphate dehydrogenase (G6PDH), lactate dehydrogenase (LDH), and citrate synthase (CS) were measured prior to (control) and after (test) FAST freezing to -75°C (see Fig. S1b for protocol). (**a, c, d**) The test aliquots were augmented with different additives administered at different concentrations (see *x* axes). The activity in the unfrozen aliquot served as control = 100%. Freezing without additive (concentration = 0) caused complete loss of the enzyme activity. Different additives protected the enzymes from loss of activity upon freezing at relatively low concentrations ($P_{50}$ is a cryoprotective concentration that allows recovering 50% of control, pre-freezing activity). (**b**) The extract aliquots augmented with proline or Ficoll were exposed to heat shocks. Neither proline nor Ficoll were able to protect the enzyme from almost complete loss of activity upon the heat shock.

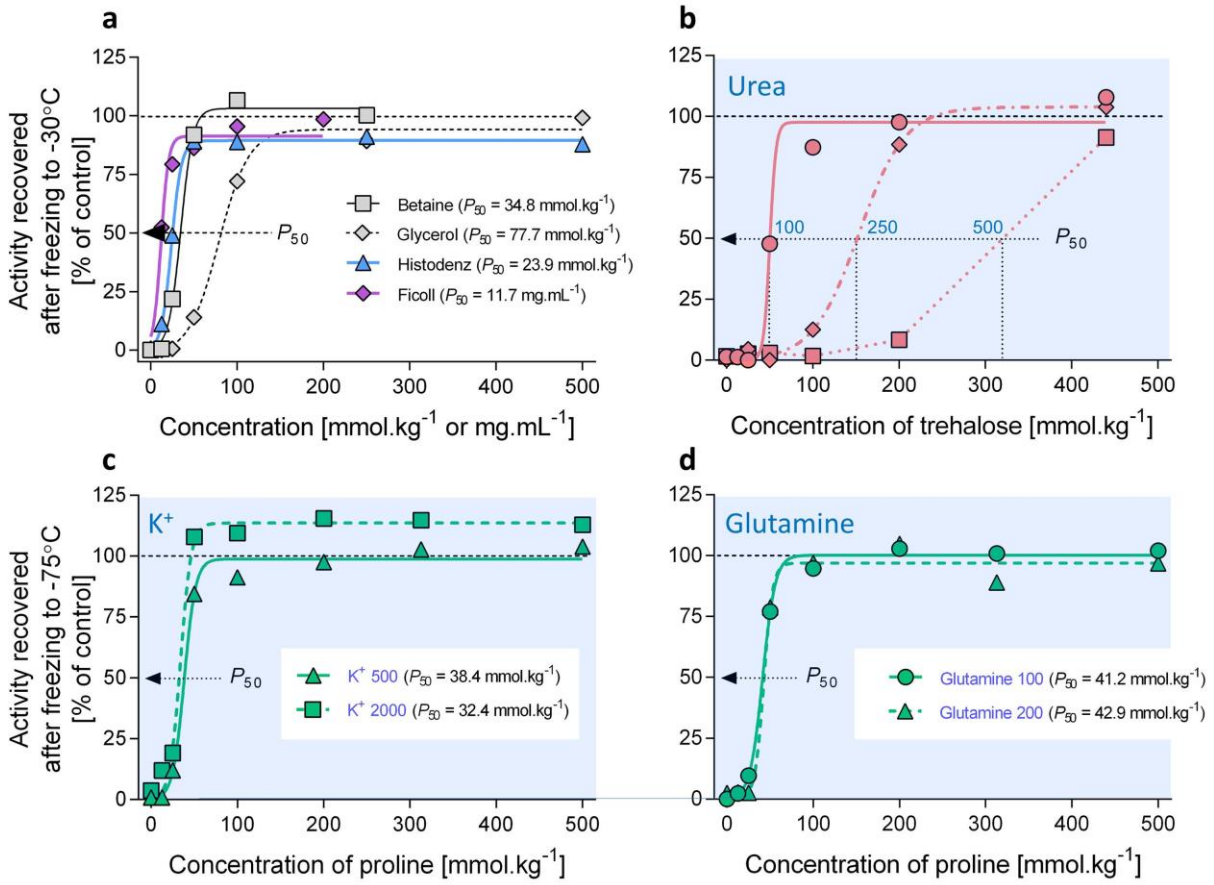


**Fig. S6: Fig. 3. Slow rate of *in vitro* freezing in the absence or presence of chemical perturbants – supplement to Fig. 3.**

The total proteins were extracted from the fat body tissue of freeze-sensitive phenotype of *Chymomyza costata* larvae and the activity of glucose 6-phosphate dehydrogenase (G6PDH) was measured prior to (control) and after (test) SLOW inoculative freezing to -30°C (see Fig. S1b for protocol). (**a**) The test aliquots were augmented with different additives administered at different concentrations (see *x* axes). The activity in the unfrozen aliquot served as control = 100%. Freezing without additive (concentration = 0) caused complete loss of enzyme activity. Different additives protected the enzyme from loss of activity upon freezing at relatively low concentrations ($P_{50}$ is a cryoprotective concentration that allows recovering 50% of control, pre-freezing activity). (**b, c, d**) Simultaneous presence of potential perturbants (urea, $K^+$, or glutamine) and cryoprotectants (trehalose or proline) during SLOW freezing either resulted in shifting the $P_{50}$ values for trehalose toward higher concentrations (b), or had negligible effect on $P_{50}$ values for proline (c, d).

**Note that** the results presented in Fig. S6b-d and also in Fig. S4b, c suggest that urea at concentrations higher than approximately 500 mmol-kg$^{-1}$ shows perturbing effects on G6PDH activity. Neither K+ nor glutamine applied at very high concentrations (close to solubility limits) showed clear perturbing effects on G6PDH activity.

**Table S2: The results of DSC analysis – part 1: Urea + Proline**

| Urea 100 +Proline | | | | | | | |
|---|---|---|---|---|---|---|---|
| Urea | Proline | water mass | melt endotherm* | delta H | frozen water (OAW fraction) | | Freeze-concentration of urea |
| $mmol.kg^{-1}$ | $mmol.kg^{-1}$ | mg | mJ | J/g | % | linear regression | $mmol.kg^{-1}$ |
| 100 | 0 | 10,00 | 3227,42 | 322,74 | 96,48 | 95,13 | 2054,1 |
| 100 | 12,5 | 9,82 | 3035,56 | 309,22 | 92,44 | 94,92 | 1970,4 |
| 100 | 25 | 10,14 | 3244,40 | 319,96 | 95,65 | 94,72 | 1893,3 |
| 100 | 50 | 10,12 | 3149,88 | 311,24 | 93,04 | 94,30 | 1755,8 |
| 100 | 100 | 10,07 | 3090,04 | 306,75 | 91,70 | 93,48 | 1533,1 |
| 100 | 200 | 10,08 | 3244,46 | 321,96 | 96,25 | 91,82 | 1222,9 |
| 100 | 313 | 9,84 | 2967,85 | 301,73 | 90,20 | 89,95 | 995,3 |
| 100 | 500 | 9,89 | 2826,57 | 285,77 | 85,43 | 86,86 | 760,9 |

| Urea 250 +Proline | | | | | | | |
|---|---|---|---|---|---|---|---|
| Urea | Proline | water mass | melt endotherm* | delta H | frozen water (OAW fraction) | | Freeze-concentration of urea |
| $mmol.kg^{-1}$ | $mmol.kg^{-1}$ | mg | mJ | J/g | % | linear regression | $mmol.kg^{-1}$ |
| 250 | 0 | 10,10 | 3167,77 | 313,69 | 93,78 | 93,85 | 4063,3 |
| 250 | 12,5 | 10,05 | 3175,45 | 315,83 | 94,42 | 93,64 | 3932,7 |
| 250 | 25 | 10,29 | 3284,67 | 319,33 | 95,46 | 93,44 | 3810,3 |
| 250 | 50 | 10,08 | 3077,01 | 305,25 | 91,26 | 93,03 | 3586,9 |
| 250 | 100 | 9,88 | 2960,98 | 299,76 | 89,61 | 92,21 | 3210,5 |
| 250 | 200 | 9,96 | 3044,94 | 305,69 | 91,39 | 90,58 | 2653,6 |
| 250 | 313 | 9,94 | 3002,32 | 301,98 | 90,28 | 88,73 | 2218,7 |
| 250 | 500 | 9,85 | 2797,80 | 284,18 | 84,96 | 85,68 | 1745,3 |

| Urea 500 +Proline | | | | | | | |
|---|---|---|---|---|---|---|---|
| Urea | Proline | water mass | melt endotherm* | delta H | frozen water (OAW fraction) | | Freeze-concentration of urea |
| $mmol.kg^{-1}$ | $mmol.kg^{-1}$ | mg | mJ | J/g | % | linear regression | $mmol.kg^{-1}$ |
| 500 | 0 | 9,81 | 3022,95 | 308,28 | 92,16 | 92,24 | 6446,4 |
| 500 | 12,5 | 9,91 | 2997,07 | 302,48 | 90,43 | 91,98 | 6236,7 |
| 500 | 25 | 10,06 | 3093,06 | 307,48 | 91,92 | 91,72 | 6040,3 |
| 500 | 50 | 9,91 | 2995,44 | 302,41 | 90,41 | 91,20 | 5682,3 |
| 500 | 100 | 9,74 | 2960,98 | 304,14 | 90,92 | 90,16 | 5080,2 |
| 500 | 200 | 9,54 | 2912,99 | 305,23 | 91,25 | 88,07 | 4191,9 |
| 500 | 313 | 9,74 | 2757,31 | 283,19 | 84,66 | 85,72 | 3500,2 |
| 500 | 500 | 9,42 | 2558,67 | 271,49 | 81,16 | 81,81 | 2749,5 |

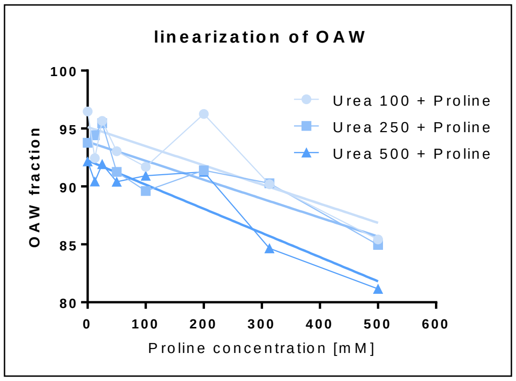


* the analysis of melt endotherm of bulk water is sensitive to the setting of baseline, which must be done manually – this causes fluctuations in the estimated OAW fraction and, consequently, in the estimated freeze-concentration of urea. In order to mitigate the effect of manually setting the baseline, we used OAW fractions interpolated from linear regressions (see figure).

**Table S2 - continues: The results of DSC analysis – part 2: Urea + Trehalose**

**Urea 100 + Trehalose**

| Urea | Trehalose | water mass | melt endotherm* | delta H | frozen water (OAW fraction) | | Freeze-concentration of urea |
|---|---|---|---|---|---|---|---|
| mmol.kg$^{-1}$ | mmol.kg$^{-1}$ | mg | mJ | J/g | % | linear regression | mmol.kg$^{-1}$ |
| 100 | 0 | 9,90 | 3180,34 | 321,23 | 96,03 | 94,85 | 1943,4 |
| 100 | 12,5 | 9,99 | 3228,96 | 323,15 | 96,61 | 94,61 | 1854,6 |
| 100 | 25 | 9,99 | 3071,75 | 307,63 | 91,97 | 94,36 | 1773,6 |
| 100 | 50 | 10,00 | 2983,69 | 298,34 | 89,19 | 93,87 | 1631,0 |
| 100 | 100 | 9,81 | 3128,40 | 318,92 | 95,34 | 92,88 | 1405,2 |
| 100 | 200 | 9,97 | 3107,89 | 311,86 | 93,23 | 90,91 | 1100,5 |
| 100 | 500 | 9,13 | 2568,51 | 281,38 | 84,12 | 85,00 | 666,7 |

**Urea 250 + Trehalose**

| Urea | Trehalose | water mass | melt endotherm* | delta H | frozen water (OAW fraction) | | Freeze-concentration of urea |
|---|---|---|---|---|---|---|---|
| mmol.kg$^{-1}$ | mmol.kg$^{-1}$ | mg | mJ | J/g | % | linear regression | mmol.kg$^{-1}$ |
| 250 | 0 | 10,08 | 3261,05 | 323,56 | 96,73 | 95,46 | 5504,8 |
| 250 | 12,5 | 9,77 | 3084,92 | 315,86 | 94,43 | 95,10 | 5099,0 |
| 250 | 25 | 9,88 | 3016,07 | 305,33 | 91,28 | 94,74 | 4749,0 |
| 250 | 50 | 9,88 | 3184,59 | 322,48 | 96,41 | 94,01 | 4175,7 |
| 250 | 100 | 9,81 | 3150,38 | 321,11 | 96,00 | 92,57 | 3363,5 |
| 250 | 200 | 9,58 | 2754,28 | 287,47 | 85,94 | 89,68 | 2421,6 |
| 250 | 500 | 9,07 | 2480,08 | 273,49 | 81,76 | 81,00 | 1316,0 |

**Urea 500 + Trehalose**

| Urea | Trehalose | water mass | melt endotherm* | delta H | frozen water (OAW fraction) | | Freeze-concentration of urea |
|---|---|---|---|---|---|---|---|
| mmol.kg$^{-1}$ | mmol.kg$^{-1}$ | mg | mJ | J/g | % | linear regression | mmol.kg$^{-1}$ |
| 500 | 0 | 10,00 | 3196,04 | 319,60 | 95,55 | 93,13 | 7273,8 |
| 500 | 12,5 | 9,91 | 3008,42 | 303,42 | 90,71 | 92,85 | 6988,2 |
| 500 | 25 | 9,89 | 3154,92 | 319,02 | 95,37 | 92,56 | 6724,3 |
| 500 | 50 | 9,88 | 3037,95 | 307,58 | 91,95 | 92,00 | 6252,0 |
| 500 | 100 | 9,76 | 2951,19 | 302,45 | 90,42 | 90,88 | 5481,9 |
| 500 | 200 | 9,60 | 2708,83 | 282,03 | 84,31 | 88,63 | 4398,3 |
| 500 | 500 | 8,76 | 2450,40 | 279,73 | 83,63 | 81,89 | 2761,1 |

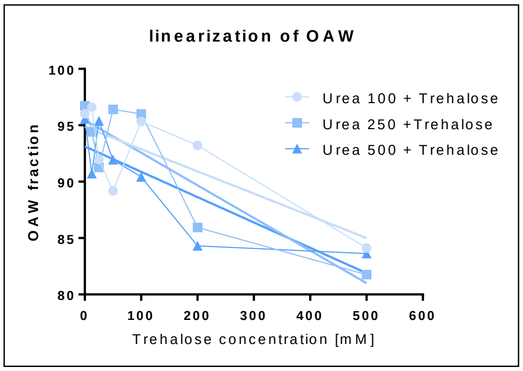


* the analysis of melt endotherm of bulk water is sensitive to the setting of baseline, which must be done manually – this causes fluctuations in the estimated OAW fraction and, consequently, in the estimated freeze-concentration of urea. In order to mitigate the effect of manually setting the baseline, we used OAW fractions interpolated from linear regressions (see figure).

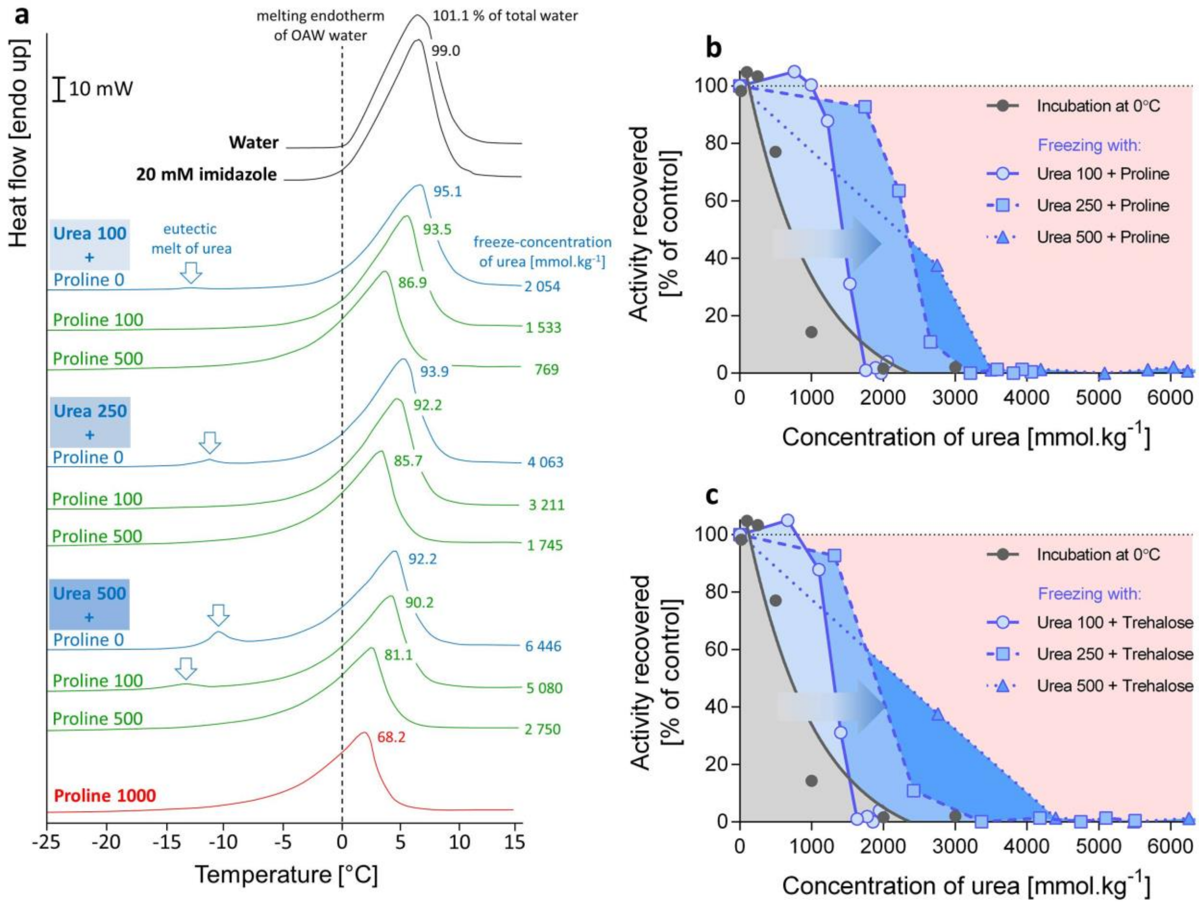


**Fig. S7: The freeze-concentration of urea caused by freezing the osmotically active fraction of water (OAW).** **(a)** Examples of DSC heating curves for the solutions of urea and proline mixed in different concentrations (values show mmol.kg$^{-1}$) dissolved in 20 mM imidazole (complete dataset is in Table S2). The area under the melt endotherm of bulk water (mJ) corresponds to the fraction of frozen, i.e. osmotically active water (OAW) in the mixture. For pure water, the OAW fraction is 100% in theory (a small deviation from theory, 101.1%, is caused mainly by manually setting the baseline). For 20 mM imidazole, the OAW fraction was close to 100%. Note that gradually increasing concentrations of urea + proline caused the gradual decrease of area under the melt endotherm (the decrease of OAW fraction) and the shift of onset of melting from 0°C (water) to subzero temperatures. The numbers close to peaks of melt endotherms show the measured OAW fraction (% of total water). The smaller is the OAW fraction, the lower is the freeze-concentration effect on urea – the estimation is shown at the end of each curve. (**b, c**) The effect of urea concentration on activity of G6PDH recovered after either the incubation of total protein extract at 0°C for 1h (taken from Fig. S4c), or after SLOW freezing to -30°C (taken from Fig. 3e for proline, and from Fig. S6b for trehalose).

**Note that** the concentrations of urea *much higher* than those set by incubation at 0°C experiment (grey area) are tolerated by G6PDH in the -30°C freezing experiment provided proline (b) or trehalose (c) are added to the solution (arrows from grey to blue areas). These results suggest that proline and trehalose not only help to decrease the freeze-concentration of urea but also directly counteract the chemical disturbance to G6PDH enzyme caused by high urea concentrations. The red area indicates that extremely high concentrations or urea are not tolerable for G6PDH even in the presence of proline or trehalose.

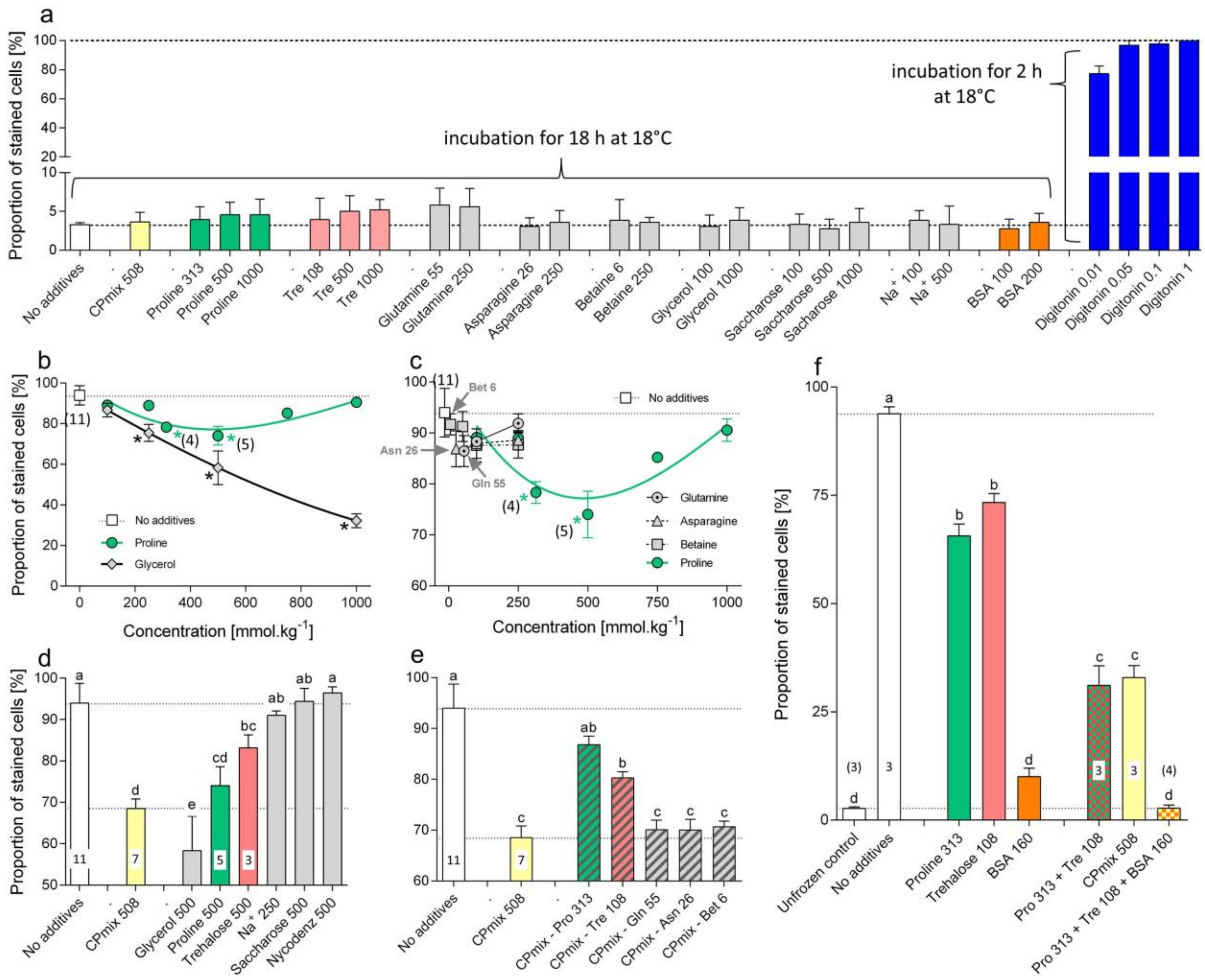


**Fig. S8: Effect of additives on plasma membrane integrity of fat body cells during incubation and freezing stress *in vitro*.**

The dissected fat body tissues of (**a-e**) freeze-sensitive, or (**f**) freeze-tolerant, phenotype larvae of *Chymomyza costata* were either (**a**) incubated at 18°C, or **(b-f**) exposed to SLOW inoculative freezing stress *in vit*ro, in Schneider's solution augmented by different additives at different concentrations (see *x* axes). The control tissues (empty squares in (b, c) or empty columns in (a, d-f) were exposed to incubation or freezing stress without additive. After incubation or melting, the integrity of fat body cell plasma membrane was assessed using Trypan Blue assay (see *y* axes). For other descriptions, see caption of Fig. 5.

**Note that:**

(**a**) Additives have no effect on membrane integrity of the cells incubated in Schneider's solution at 18°C, while Digitonin at low concentrations permeabilize the membrane for Trypan Blue.

(**b, c**) Glycerol is more effective in membrane integrity protection than proline, while glutamine, asparagine and betaine are ineffective (data for proline are shown for comparison).

(**d**) Different additives applied in same osmotic concentration showed different ability to protect membrane integrity.

(**e**) Excluding proline or trehalose from CPmix caused partial loss of the ability of reduced-CPmix to protect membrane integrity.

(**f**) Complete rescue of membrane integrity upon *in vitro* freezing stress was achieved in freeze-tolerant phenotype larvae using mixture of proline, trehalose and BSA.

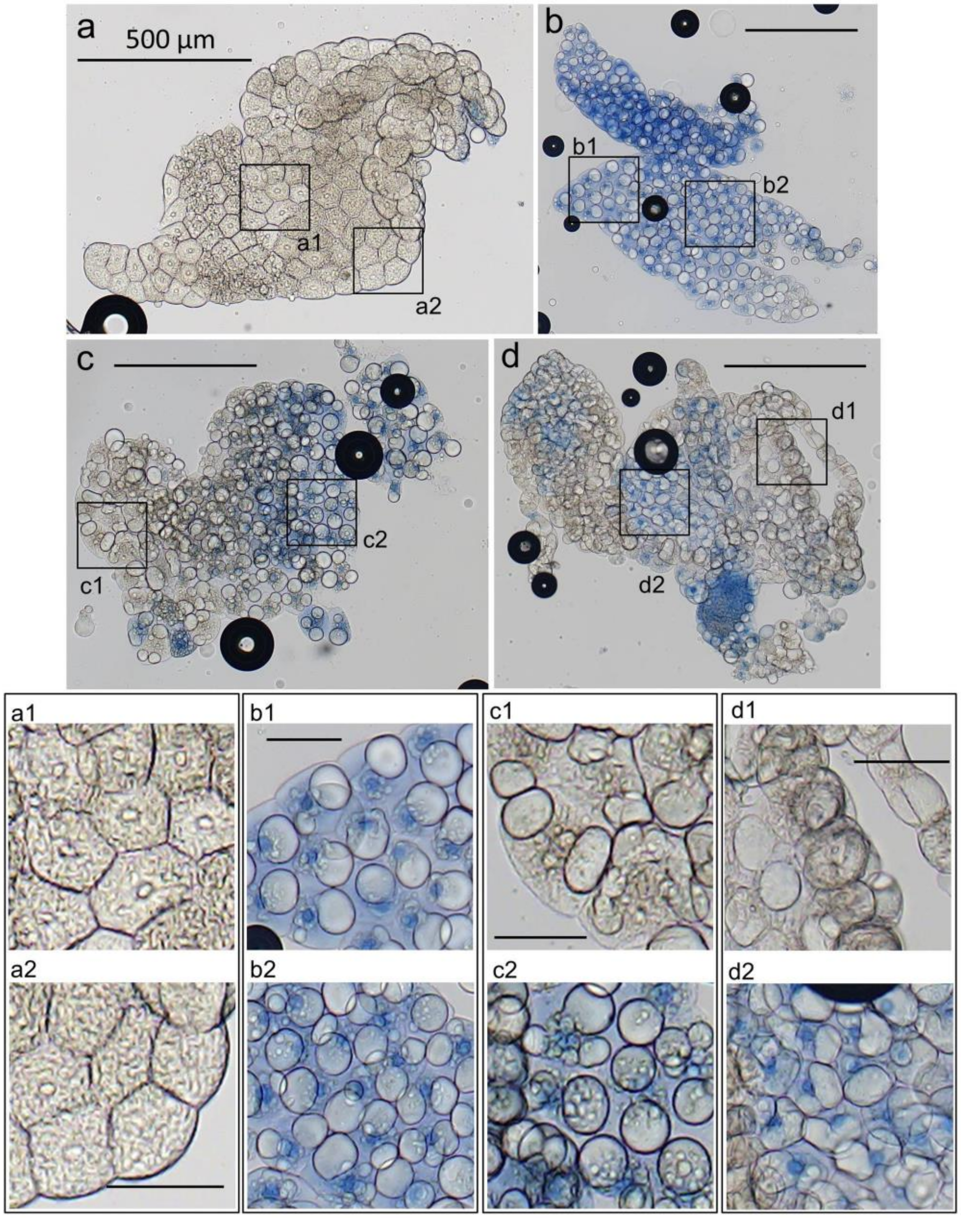


**Fig. S9: Examples of Trypan Blue staining of fat body cells.**
The fat body tissue was dissected from freeze-sensitive phenotype of *Chymomyza costata* larvae. Controls (**a**) were not exposed to freezing stress and incubated directly in Trypan Blue. The test tissues were first exposed to SLOW inoculative freezing to -30°C and incubated in Trypan Blue after melting. The cells with compromised membrane integrity stain blue. The test tissues were frozen in: (**b**) plain Schneider's *Drosophila* medium; or (**c**) Schneider's medium augmented with CPmix 508 (mixture of proline, trehalose, glutamine, asparagine, and betaine in total concentration of 508 mmol.kg$^{-1}$); or (**d**) Schneider's medium augmented with BSA, 100 mg.mL$^{-1}$. All scale bars in (a-d) are 500 µm. The squares delimit two areas in each micrograph that are shown enlarged below.

**Note that** each piece of tissue shows compact areas with relatively intact cells (no staining), while the cells are stained with Trypan Blue in other areas. All frozen tissues show the characteristic coalescence of initially small lipid droplets (a) into several or single large lipid droplet (b-d) (17).

#### SI Supplementary Discussion

**Exact quotations of highly-cited literary sources:**

- Storey and Storey, 1988 (18), p. 43 (742 citations in Google Scholar): " ... the presence of polyols in freeze-tolerant insects may be key to preserving homeostasis in the frozen state by stabilizing individual enzymes/proteins as well as macromolecular interactions ... against the potential denaturing effects of subzero temperatures and low water content."

- Lee, 2010 (19), p. 17 (389 citations in Google Scholar):" Trehalose and glycerol are commonly produced by insects in response to osmotic challenge due to desiccation and freezing. These compounds serve to stabilize proteins and protect cell membranes."

- Teets and Denlinger, 2013 (20), p. 3 (266 citations in Google Scholar): "The most common cryoprotectants observed in overwintering insects are low molecular weight sugar alcohols such as glycerol, sorbitol and inositol. The functions of these compounds include stabilizing membranes and proteins ..."

- Toxopeus and Sinclair, 2018 (21) (91 citations in Google Scholar), p 9: "Increased osmotic pressure associated with freezing (i.e. freeze concentration) might destabilise proteins and damage cell membranes, causing cell death." p. 13: " ... freeze-tolerant insects must counteract the destabilising effect of dehydration on membranes (including organelle and vesicle membranes) and proteins. These macromolecules can be stabilised by direct or indirect (*via* the hydration shell) interaction with low molecular weight metabolites such as trehalose, proline, and other amino acids."

- The numbers of citations are according to Google Scholar (https://scholar.google.com/) taken on 15 June 2022.

**Do proteins in general need stabilization by CPs during freezing stress?**
There are several reasons for cautiousness when extending the validity of our hypothesis that the insect *soluble enzymes* are not the primary targets of freezing injury to *all proteins* in general:

First, dissociation of multimeric proteins into subunits was observed to occur in many enzymes at temperatures close to 0°C (22). Similarly, polymeric protein structures such as microtubules, actin filaments, or collagen fibers depolymerize already upon mild cold exposures (23-25). Such dissociation of quaternary structures represents a first step toward protein denaturation and is usually associated with loss of protein function. Nevertheless, spontaneous re-association of mild-cold-dissociated enzymes and polymeric proteins was observed upon incubation at physiological temperatures. Most of the enzymes assessed in this study are dimers or tetramers and their dissociation upon freezing stress *in vivo* (if it occurred at all) must have been fully reversible upon melting and re-warming because no loss of activity was observed. The re-association of larger complexes after severe cold and freeze-dehydration stress, however, might be more problematic as the osmotic shrinkage of cells causes loss of cytoplasmic organization, extensive displacements or even disintegration of organelles and cytoskeletal structures (10, 17, 26).

Second, severe cold alone (typically around -20°C) is known to cause cold-denaturation of proteins, i.e. unfolding of globular tertiary structure and exposing hydrophobic core domains (22, 27). The occurrence of protein denaturation is theoretically even more probable after freezing due to loss of bulk water and increasing concentrations of chemical perturbants of native protein structure such as protons, metal ions, or urea (28-30). Again, the large polymeric protein complexes might be more sensitive than soluble enzymes to irreversible aggregation of denatured subunits via their exposed hydrophobic domains upon severe freezing stress. Accordingly, the freeze-thaw induced denaturation and aggregation of myofibrilar proteins was often observed in fish muscle (31, 32). We observed irreversible loss of microtubular structure to occur in fat body cells of frozen-thawed larvae of *C. costata* – however, without any apparent loss of larval viability (17).

Third, the thermodynamic theory of protein denaturation by cold and freezing stress, and its prevention by CP-mediated stabilization, has been well established during 80-ties of the last century (33-35) and is accepted until recently (36, 37). That some insect proteins really do denature upon cold and freezing stress *in vivo* is indirectly supported by the observed upregulation of inducible heat shock proteins' expression after the stress (38, 39), as well as by the impaired cold tolerance of insects that had this upregulation response prevented by RNAi targeted against expression of *hsp70* gene (40). We found that the *in vitro* loss of activity of G6PDH caused by heat stress (+45°C or +65°C for 1h) was not preventable by proline or Ficoll, while the loss of activity caused by freezing stress (-75°C for 1h) was fully preventable by low concentrations of proline and Ficoll. This result suggests that the nature or extent of the injury to proteins differ between heat and freezing stresses.

Fourth, the accumulated CPs may compensate for the denaturing effects of high concentrations of chemical perturbants (41-43). The concentrations of protein structure denaturants such as urea, arginine, lysine, or inorganic salts may, in theory, exceed toxic limits due to freeze dehydration of biological solutions (29). It has been shown, for instance, that elasmobranch fish accumulating high concentrations of urea (400 mM) protect their pyruvate kinase and ribonuclease against denaturing effects of urea by a mixture of compensating solutes containing TMAO, sarcosine, β-alanine, and betaine (44, 45). Accordingly, we observed that G6PDH activity is protected during freezing stress by proline or trehalose against perturbing effects of urea concentrations exceeding 500 $mmol.kg^{-1}$. Such high concentrations of urea, however, are unlikely to occur in larvae. We have not measured the concentrations of urea in larvae of *C. costata*; nevertheless, data for larvae of *Drosophila melanogaster* show hemolymph concentrations below 10 $mmol.l^{-1}$ (46).

Fifth, we have studied the acute effects caused by single freeze-thaw cycle, while the overwintering insects are often exposed to long-term freezing and/or repeated freeze-thaw cycles in the field (47). The increasing time of frozen-storage and number of freeze-thaw cycles are well known to aggravate the irreversible denaturation of fish myofibrilar proteins (32).

Collectively, for the reasons specified above, we keep open the question on stability of *proteins in general* upon freezing stress. The stability of large polymeric protein complexes, nucleoprotein multimeric complexes such as ribosomes, or membrane-embedded protein complexes should be attested specifically in independent experiments.

#### SI References


1. Riihimaa AJ & Kimura MT (1988) A mutant strain of *Chymomyza costata* (Diptera: Drosophilidae) insensitive to diapause-inducing action of photoperiod. *Physiol Entomol.* 13(4):441-445.
2. Kostal V, Noguchi H, Shimada K, & Hayakawa Y (1998) Developmental changes in dopamine levels in larvae of the fly Chymomyza costata: comparison between wild-type and mutant-nondiapause strains. *J. Insect Physiol.* 44(7-8):605-614.
3. Kučera L*, et al.* (2022) A mixture of innate cryoprotectants is key for freeze tolerance and cryopreservation of a drosophilid fly larva. *J. Exp. Biol.* 225(8):jeb243934.
4. Rozsypal J, Moos M, Šimek P, & Koštál V (2018) Thermal analysis of ice and glass transitions in insects that do and do not survive freezing. *J. Exp. Biol.* 221:170464.
5. Koštál V*, et al.* (2011) Long-term cold acclimation extends survival time at 0°C and modifies the metabolomic profiles of the larvae of the fruit fly *Drosophila melanogaster*. *PLoS One* 6(9):e25025.
6. Findsen A, Andersen JL, Calderon S, & Overgaard J (2013) Rapid cold hardening improves recovery of ion homeostasis and chill coma recovery time in the migratory locust, *Locusta migratoria*. *J. Exp. Biol.* 216(9):1630-1637.
7. Koštál V, Zahradníčková H, & Šimek P (2011) Hyperprolinemic larvae of the drosophilid fly, Chymomyza costata, survive cryopreservation in liquid nitrogen. *Proc. Natl. Acad. Sci. U.S.A.* 108(32):13041-13046.
8. Smith Pe*, et al.* (1985) Measurement of protein using bicinchoninic acid. *Anal. Biochem.* 150(1):76-85.
9. Storey KB, Keefe D, Kourtz L, & Storey JM (1991) Glucose-6-phosphate dehydrogenase in cold hardy insects: kinetic properties, freezing stabilization, and control of hexose monophosphate shunt activity. *Insect Biochem.* 21(2):157-164.
10. Štětina T, Des Marteaux L, & Koštál V (2020) Insect mitochondria as targets of freezing-induced injury. *Proc. R. Soc. B* 287(1931):20201273.
11. Carpenter JF & Crowe JH (1988) The mechanism of cryoprotection of proteins by solutes. *Cryobiology* 25(3):244-255.
12. Laughton AM & Siva-Jothy MT (2011) A standardised protocol for measuring phenoloxidase and prophenoloxidase in the honey bee, Apis mellifera. *Apidologie* 42(2):140-149.
13. Koštál V, Tollarova M, & Šula J (2004) Adjustments of the enzymatic complement for polyol biosynthesis and accumulation in diapausing cold-acclimated adults of Pyrrhocoris apterus. *J. Insect Physiol.* 50(4):303-313.
14. Storey KB & Storey JM (2012) Insect cold hardiness: metabolic, gene, and protein adaptation. *Can. J. Zool.* 90(4):456-475.
15. Strober W (2015) Trypan blue exclusion test of cell viability. *Curr. Protoc. Immunol.* 111(1):A3. B. 1-A3. B. 3.
16. Fiskum G (1985) Intracellular levels and distribution of Ca2+ in digitonin-permeabilized cells. *Cell Calcium* 6(1-2):25-37.
17. Des Marteaux LE, Štětina T, & Koštál V (2018) Insect fat body cell morphology and response to cold stress is modulated by acclimation. *J. Exp. Biol.* 221(21):jeb189647.
18. Storey KB & Storey JM (1988) Freeze tolerance in animals. *Physiol. Rev.* 68(1):27-84.

19. Lee REJ (2010) A primer on insect cold-tolerance. *Low Temperature Biology of Insects*, eds Denlinger DL & Lee REJ (Cambridge University Press).
20. Teets NM & Denlinger DL (2013) Physiological mechanisms of seasonal and rapid cold-hardening in insects. *Physiol. Entomol.* 38(2):105-116.
21. Toxopeus J & Sinclair BJ (2018) Mechanisms underlying insect freeze tolerance. *Biol. Rev.* 93:1891-1914.
22. Privalov PL (1990) Cold denaturation of protein. *Crit. Rev. Biochem. Mol. Biol.* 25(4):281-306.
23. Cottam DM*, et al.* (2006) Non-centrosomal microtubule-organising centres in cold-treated cultured *Drosophila* cells. *Cell Motil. Cytoskeleton* 63(2):88-100.
24. Belous A (1992) The role of regulatory systems modulating the state of cytoskeletal proteins under the temperature and osmotic effects. *Probl. Cryobiol.* 4:3-14.
25. Fessler J (1960) Some properties of neutral-salt-soluble collagen. 2. *Biochem. J.* 76(3):463.
26. Des Marteaux LE, Stinziano JR, & Sinclair BJ (2018) Effects of cold acclimation on rectal macromorphology, ultrastructure, and cytoskeletal stability in *Gryllus pennsylvanicus* crickets. *J. Insect Physiol.* 104:15-24.
27. Dias CL*, et al.* (2010) The hydrophobic effect and its role in cold denaturation. *Cryobiology* 60(1):91-99.
28. Bennion BJ & Daggett V (2003) The molecular basis for the chemical denaturation of proteins by urea. *Proc. Natl. Acad. Sci. U.S.A.* 100(9):5142-5147.
29. Timasheff SN (1993) The control of protein stability and association by weak interactions with water: how do solvents affect these processes? *Annu. Rev. Biophys. Biomol. Struct.* 22(1):67-97.
30. Hamada H, Arakawa T, & Shiraki K (2009) Effect of additives on protein aggregation. *Curr. Pharm. Biotechnol.* 10(4):400-407.
31. Zhang Y, Puolanne E, & Ertbjerg P (2021) Mimicking myofibrillar protein denaturation in frozen-thawed meat: Effect of pH at high ionic strength. *Food Chem.* 338:128017.
32. Sikorski ZE, Olley J, Kostuch S, & Olcott HS (1976) Protein changes in frozen fish. *Crit. Rev. Food Sci. Nutr.* 8(1):97-129.
33. Gekko K & Timasheff SN (1981) Mechanism of protein stabilization by glycerol: preferential hydration in glycerol-water mixtures. *Biochemistry* 20(16):4667-4676.
34. Arakawa T & Timasheff SN (1982) Stabilization of protein structure by sugars. *Biochemistry* 21(25):6536-6544.
35. Arakawa T & Timasheff S (1985) The stabilization of proteins by osmolytes. *Biophys. J.* 47(3):411-414.
36. Kim NA, Thapa R, & Jeong SH (2018) Preferential exclusion mechanism by carbohydrates on protein stabilization using thermodynamic evaluation. *Int. J. Biol. Macromol.* 109:311-322.
37. Arsiccio A & Pisano R (2017) Stability of proteins in carbohydrates and other additives during freezing: the human growth hormone as a case study. *J. Phys. Chem. B* 121(37):8652-8660.
38. Sinclair B, Gibbs A, & Roberts S (2007) Gene transcription during exposure to, and recovery from, cold and desiccation stress in *Drosophila melanogaster*. *Insect Mol. Biol.* 16(4):435-443.
39. Teets NM*, et al.* (2019) Changes in energy reserves and gene expression elicited by freezing and supercooling in the Antarctic midge, Belgica antarctica. *Insects* 11(1):18.
40. Rinehart JP*, et al.* (2007) Up-regulation of heat shock proteins is essential for cold survival during insect diapause. *Proc. Natl. Acad. Sci. U.S.A.* 104(27):11130-11137.
41. Somero G (1986) Protons, osmolytes, and fitness of internal milieu for protein function. *Am. J. Physiol. Regul. Integr. Comp. Physiol.* 251(2):R197-R213.
42. Yancey PH (2005) Organic osmolytes as compatible, metabolic and counteracting cytoprotectants in high osmolarity and other stresses. *J. Exp. Biol.* 208(15):2819-2830.

43. Zachariassen KE, Kristiansen E, & Pedersen SA (2004) Inorganic ions in cold-hardiness. *Cryobiology* 48(2):126-133.
44. Yancey PH & Somero GN (1979) Counteraction of urea destabilization of protein structure by methylamine osmoregulatory compounds of elasmobranch fishes. *Biochem. J.* 183(2):317-323.
45. Yancey PH & Somero GN (1980) Methylamine osmoregulatory solutes of elasmobranch fishes counteract urea inhibition of enzymes. *J. Exp. Zool.* 212(2):205-213.
46. Etienne R, Fortunat K, & Pierce V (2001) Mechanisms of urea tolerance in urea-adapted populations of Drosophila melanogaster. *J. Exp. Biol.* 204(15):2699-2707.
47. Marshall KE & Sinclair BJ (2015) The relative importance of number, duration and intensity of cold stress events in determining survival and energetics of an overwintering insect. *Funct. Ecol.* 29(3):357-366.
